# Testing Thirty Years of Hypotheses: Ancient Trout–Salmon Introgression and Limited Pre-Anthropocene Gene Flow Between Rainbow and Cutthroat Trout Shaped Pacific Salmonid Evolution

**DOI:** 10.64898/2026.10.12.758086

**Authors:** Ethan R. Tolman, Harrison Pardue, Gracie Castillo, Anna Eichert, James Holt, Dick Jordan, Emma Redman, Liam Williams, Anna Himler, Christoph Walser, Christopher D. Beatty, Anton Suvorov

## Abstract

Introgression is the transfer of genetic material between species or populations through hybridization and backcrossing. While often deleterious or neutral in nature, introgression can also have beneficial consequences for the recipient lineage. *Oncorhynchus* is a genus of ray-finned fish (family Salmonidae) composed of nine species of Pacific trout and eight species of Pacific salmon, in which deep introgression has long been proposed to explain patterns of gene-tree-species-tree discordance, and in which hybridization is extensively observed between introduced and endemic species. In this study, we identified well-supported introgression between the ancestral lineage of Pacific trout and the ancestor of Chinook and coho salmon, a long-posited but previously untested hypothesis. While other salmon lineages (pink, chum, sockeye, and masu salmon) have been sympatric with trout for extended periods, providing an opportunity for hybridization to occur, we found no evidence of introgression between trout and these other salmon lineages. Our findings suggest mito-nuclear incompatibilities serve as a possible barrier to gene flow between the clade comprised of pink, chum, and sockeye salmon and all other lineages in *Oncorhynchus*. Within Pacific trout, we detected no historical introgression between cutthroat and rainbow trout lineages, suggesting that the extensive hybridization observed between them today reflects recent, primarily human-mediated contact rather than long-term gene flow. These findings show that ancestral introgression has shaped the evolutionary history of *Oncorhynchus* and should be accounted for in future comparative phylogenetic studies of the genus.

## Introduction

### Overview of introgression and gene flow

Introgression is a key evolutionary process observed in many lineages across the Tree of Life with a genetically homogenizing effect (Suarez-Gonzalez et al. 2018; Qian et al. 2023). Widespread signatures of introgression have been observed in a variety of taxa such as plants (Suarez-Gonzalez et al. 2018), vertebrates (Zinner et al. 2009; Hallström and Janke 2010; Hohenlohe et al. 2013; Bernal et al. 2017; Kumar et al. 2017; Singhal et al. 2021; Koller et al. 2022; Reilly et al. 2022; Qian et al. 2023), and invertebrates (Edelman et al. 2019; Suvorov et al. 2022a; Qian et al. 2023; Wu et al. 2024).

Introgression has traditionally been thought to most commonly result in neutral or maladaptive consequences via outbreeding depression, accumulation of deleterious mutations, and genetic swamping (Kleindorfer et al. 2014; Adavoudi and Pilot 2021). While this remains true and well-documented, recent advances in genomic tools have helped to identify a growing number of lineages showing historical adaptive introgression across the Tree of Life (reviewed in Arnold and Kunte 2017; Horta et al. 2025), including, but not limited to, cichlids (Meier et al. 2017), butterflies (Edelman et al. 2019), corals (Wu et al. 2024), and hominids (Sankararaman et al. 2014; Vernot and Akey 2014; Huerta-Sánchez et al. 2014; Koller et al. 2022; Reilly et al. 2022; Iasi et al. 2024).

Geographically sympatric lineages have greater opportunity to hybridize, and introgression is typically more extensive between sympatric than allopatric lineages (Martin et al. 2013; Waters et al. 2023). Yet, geographic overlap is not always predictive of the prevalence or strength of introgression (Hamlin et al. 2020). In fact, prolonged sympatry may also strengthen barriers to gene flow, through reinforcement of prezygotic isolation (Coyne and Orr 1989, 1997), spatial segregation of breeding habitat (Buehrens et al. 2013), chromosomal rearrangements which impose barriers by suppressing recombination or reducing hybrid fertility (Rieseberg 2001; Zhang et al. 2021), or mitonuclear incompatibilities that reduce fitness of hybrid offspring (Burton et al. 2013; Moran et al. 2024). Although genomic data have revealed introgression to be widespread (Mallet et al. 2016; Taylor and Larson 2019), the factors governing when sympatric lineages exchange genes, and when they do not, remain poorly understood (Hamlin et al. 2020; Moran et al. 2021; Sianta et al. 2024).

### Sympatric introgression and reproductive isolation in the fish genus Oncorhynchus

With high overlap in species ranges (Fig. 1A,B; Crawford and Muir 2008), the presence or absence of introgression in the ray-finned fish (Actinopterygii) genus *Oncorhynchus* (Salmonidae; named after a Russian term meaning “hooknose”, which refers to the hooked upper jaw males develop during mating [Page et al. 2023]) is an intriguing case study in introgression, and a lack thereof, in sympatric lineages.

**Figure 1.**
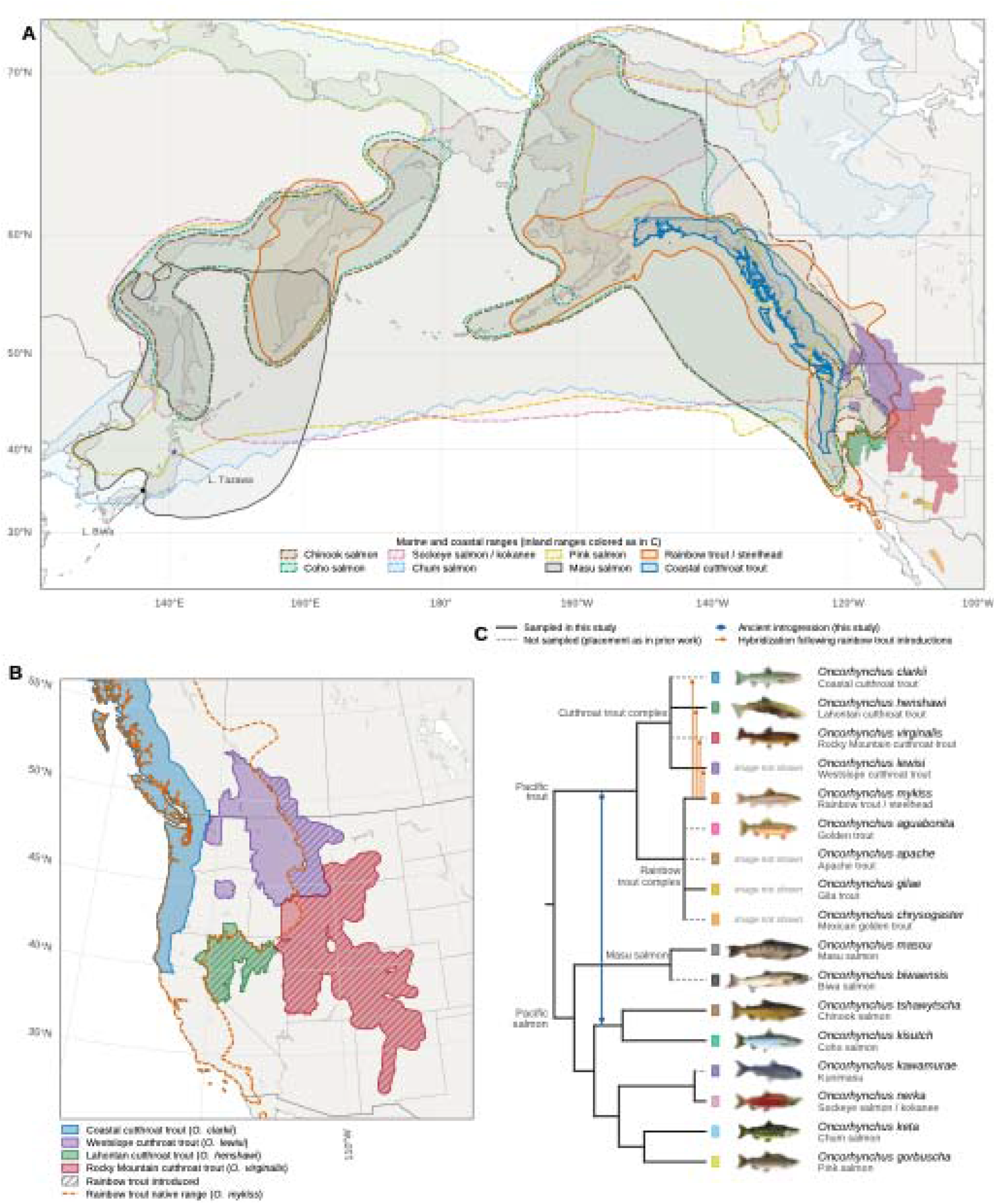
Native ranges and relationships of Oncorhynchus. A) Native distributions of extant Oncorhynchus. Marine and coastal ranges are outlined and inland ranges are colored as in C; markers show the lake endemics O. kawamurae (Lake Tazawa) and O. biwaensis (Lake Biwa). Ranges follow Crawford and Muir (2008), with cutthroat trout ranges delimited by drainage (Behnke et al. 2002; US Geological Survey 2018). B) Native ranges of the four cutthroat trout species (Page et al. 2023) and of rainbow trout (O. mykiss; dashed line). Hatching marks the parts of each cutthroat trout range, outside the native range of rainbow trout, that lie in states and provinces into which rainbow trout have been introduced (Crawford and Muir 2008). C) Hypothesized relationships within Oncorhynchus (Crête-Lafrenière et al. 2012; Loxterman and Keeley 2012; Muto et al. 2013; Campbell et al. 2023; Fujioka et al. 2025). Branch lengths are not scaled to time; dashed branches indicate species not sampled in this study. The blue arrow shows previously hypothesized introgression between the ancestor of Pacific trout and the ancestor of O. kisutch and O. tshawytscha (Utter and Allendorf 1994), and orange arrows show areas of potential hybridization of rainbow trout with cutthroat trout lineages following rainbow trout introductions. Illustrations (backgrounds removed; some mirrored) are from Wikimedia Commons: O. clarkii, A. Hoen & Co.; O. henshawi, US Fish and Wildlife Service (USFWS) Nevada Fish and Wildlife Office; O. virginalis, Birds and Nature 15 (1904); O. mykiss, D. Raver, USFWS; O. aguabonita, C. B. Hudson (1906 plate, Golden Trout Creek); O. tshawytscha, US Government (Lake Washington Ship Canal fish ladder pamphlet); O. kisutch, O. nerka, O. keta, and O. gorbuscha, T. Knepp, USFWS (all public domain); O. masou, Apple2000 (CC BY 3.0); O. biwaensis, OAS (CC BY-SA 3.0); O. kawamurae, High speed railway (CC BY-SA 4.0).

### Overview of the genus Oncorhynchus

*Oncorhynchus* contains two clades, Pacific salmon and Pacific trout (Fig. 1C). Although species boundaries in *Oncorhynchus* are still debated (Behnke et al. 2002; Markle 2018; Bestgen et al. 2019; Page et al. 2023; Christensen et al. 2025), the genus is, at present, generally considered to be composed of eight Pacific salmon (*O. tshawytscha*, Chinook; *O. kisutch*, coho; *O. nerka*, sockeye; *O. gorbuscha*, pink; *O. keta*, chum; *O. masou*, masu/cherry salmon; *O. biwaensis*, Biwa salmon; and *O. kawamurae*, kunimasu/black kokanee [note *O. kawamurae* is commonly referred to as a “trout” but it is most closely related to the salmon *O. nerka*]) and nine Pacific trout (*O. mykiss*, rainbow trout; *O. clarkii*, coastal cutthroat trout; *O. lewisi*, westslope cutthroat trout; *O. henshawi*, Lahontan cutthroat trout; *O. virginalis*, Rocky Mountain cutthroat trout; *O. gilae*, Gila trout; *O. apache*, Apache trout; *O. chrysogaster*, Mexican golden trout; and *O. aguabonita*, golden trout). Collectively these taxa occupy the North Pacific rim and its adjacent interior drainages: the anadromous Pacific salmon and *O. mykiss* range along both sides of the North Pacific, from Japan, Korea, and the Russian Far East to western North America (Alaska south to California, and to Baja California in *O. mykiss*), with *O. masou* and the Japanese endemics *O. biwaensis* and *O. kawamurae* restricted to Asia (Fig. 1A; Crawford and Muir 2008; Fujioka et al. 2025). The remaining Pacific trout are confined to western North America and northwestern Mexico, where the cutthroat lineages (*O. clarkii* coastally from Alaska to northern California; *O. lewisi*, *O. henshawi*, and *O. virginalis* across interior Columbia, Great Basin, and Rocky Mountain drainages) give way to narrowly endemic southern forms — *O. aguabonita* in the Kern River basin of the southern Sierra Nevada, *O. gilae* and *O. apache* in the Gila and Salt river headwaters of Arizona and New Mexico, and *O. chrysogaster* in the Sierra Madre Occidental (Crawford and Muir 2008).

Estimated to have diverged from its sister lineage *Salvelinus* (which includes present-day chars such as Arctic char, brook trout, and lake trout) in the Early Miocene (∼20-25 Ma; Lecaudey et al., 2018), the radiation of *Oncorhynchus* seemingly aligns with geological and climatic dynamics including tectonic activity and glaciation cycles (Montgomery 2000; Waples et al. 2008; Pitman et al. 2020). The genus continued to diversify in the late Pliocene and early Pleistocene via repeating glaciation cycles, which carved new habitat into North America and are hypothesized to have driven the divergence of *Oncorhynchus* lineages (Pitman et al. 2020). The collective native range of *Oncorhynchus* formed from these events incorporates the entirety of the Pacific Rim, including the inland drainages and river systems of Asia and North America (Crawford and Muir 2008). This genus has been introduced globally due to anthropogenic influence, and several species, notably rainbow trout (*Oncorhynchus mykiss*), have successfully established reproducing populations as invasive species (Crawford and Muir 2008).

### Previous study of introgression in Oncorhynchus

Introgression has long been hypothesized as an explanation for the variability in species tree topologies and divergence times previously recovered in this genus (Stearley and Smith 1993; Utter and Allendorf 1994). Specifically, the ancestor of coho (*O. kisutch*) and Chinook (*O. tshawytscha*) salmon has been hypothesized to have introgressed with the ancestor of Pacific trout based on similar mitochondrial gene sequences between these two lineages relative to nuclear loci (Fig. 1C; Utter and Allendorf 1994). Observations of gene-tree species-tree discordance in small nuclear locus datasets have been suggested to support the hypothesis of ancestral introgression between rainbow (*O. mykiss*) and cutthroat (*O. lewisi*, *O. clarkii*, and *O. virginalis*) trout lineages (Campton and Utter 1985; Allendorf and Leary 1988). This hypothesis is supported by the observation of multiple recent hybridization events between cutthroat trout lineages and human-introduced populations of *O. mykiss* in the historical range of cutthroat trout (Hitt et al. 2003; Rasmussen et al. 2010; Hohenlohe et al. 2011, 2013; Muhlfeld et al. 2014). However, the hypotheses discussed here have not been tested within a phylogenomic framework leveraging whole-genome data.

Not every pair of historically sympatric lineages in *Oncorhynchus* is hypothesized to have introgressed. Given species of *Oncorhynchus* with overlapping ranges, what could prevent introgression? Possibilities could include decreased hybrid fitness through processes such as mitochondrial-nuclear genome incompatibilities (Moran et al. 2024), post-zygotic barriers such as chromosomal inversions (Noor et al. 2001; Feder and Nosil 2009); perhaps species ranges have only recently come to overlap, or rampant introgression across *Oncorhynchus* is not detectable using only limited datasets with few loci that have been primarily used to infer the evolutionary history of this genus.

### Cultural and Economic Consequences of Introgression in Oncorhynchus

The importance of the identification of introgression in *Oncorhynchus* extends beyond value to the study of evolutionary processes. The identification of genetic introgression and the resulting fitness impacts is a compelling topic for stakeholders with interest in the long term survival of these species, such as regulatory agencies, industry for which *Oncorhynchus* provides important economic value and to Indigenous nations for which *Oncorhynchus* holds cultural importance (Hargrove et al. 2024).

The economic impact of *Oncorhynchus* is immense. Between 2012 and 2015, species of *Oncorhynchus* inhabiting the Salish Sea (which spans Washington State, USA and British Columbia, Canada) alone have generated an average of $1.1 billion annually along with thousands of jobs in both the United States and British Columbia (US Environmental Protection Agency 2013). Because of their anadromous life history, many species of *Oncorhynchus* play the ecologically crucial role of nutrient transport from the Pacific Ocean inland (Zhivotovsky 2015), a highly valuable ecosystem service.

*Oncorhynchus* are also of great cultural importance (Criddle and Shimizu 2014; Columbia River Inter-Tribal Fish Commission n.d.). In East Asia, Kamchatka Peninsula (Russia) salmon have long underpinned the subsistence economy of the Indigenous Koryak and remain a traditional food (Kasten 2012). Salmon in Japan are indispensable to the diet, material culture, and ritual life of the Indigenous Ainu of Hokkaido (Iwasaki-Goodman and Nomoto 2001). Japan also maintains one of the largest salmon hatchery programs in the world, centered on chum salmon (*O. keta*), with Hokkaido hatcheries alone releasing roughly one billion fry each year (Kitada 2014; Waycott 2017).

Five species *of* Oncorhynchus are found in the historical homeland of the Nez Perce, a self-governing Native American nation (Welch 2025), in North America (southern Columbia Plateau across what are now Idaho, northeastern Oregon, and southeastern Washington, centered on the Snake, Salmon and Clearwater River drainages). The following, written by one of the authors (J.H.), a member of the Nez Perce nation, reflects the importance of Pacific salmon and trout to the Nez Perce. Salmon are among the most honored of beings to the Nez Perce people. The reason for this reverence began eons ago. During the time before human beings, Creator asked all the living beings what they would provide for the humans who would be coming to the land. The first to come forward was Salmon. Salmon stated he would give all of himself to human beings. Due to Salmon being first in line and offering so much to humans, they are highly honored in ceremonies. To this day the Nez Perce people honor salmon in the exact way their ancestors have done since time immemorial. Similarly, Steelhead (anadromous rainbow trout) stated he would give his body to human beings. Further, Steelhead went on to state that humans would use their skin for certain things. To this day, Salmon and Steelhead travel to the ocean to grow big and strong before returning to spawn. They grow their strength to fulfill their relationship with human beings and to persist for their future generations.

Because of widespread introduction of rainbow trout into the ranges of cutthroat trout (Fig. 1B; Hitt et al. 2003; Rasmussen et al. 2010; Hohenlohe et al. 2011, 2013; Muhlfeld et al. 2014), the inference of gene-flow patterns between *O. mykiss* and cutthroat trout lineages prior to the human-mediated introduction of *O. mykiss* has perhaps the most urgent conservation and management implications within the genus. The existence of a historical introgression dynamic between these lineages is directly relevant to management decisions such as protection for hybridized lineages, and whether genetically “pure” populations of cutthroat trout should be prevented from hybridizing to ensure the long term survival of these lineages (Allendorf et al. 2001; Bohling 2016; Wayne and Shaffer 2016; vonHoldt et al. 2018; Quilodrán et al. 2020); this has been specifically debated with regard to *O. lewisi* within the cutthroat trout species complex (Allendorf et al. 2004; Campton and Kaeding 2005). The identification of introgression predating human introduction of rainbow trout into cutthroat trout habitat would reveal whether humans are facilitating a historical norm, or creating an entirely new dynamic.

### Objectives

Despite extensive study of individual species within *Oncorhynchus* (Berthelot et al. 2014; Colín et al. 2023; Rondeau et al. 2023a; Christensen et al. 2025), phylogenomic study across the whole genus is lacking. The most recent genus-level phylogenetic inferences were published in the 1990s to the 2000s, most of which primarily used few molecular markers (typically < 10 loci) for analysis (Stearley and Smith 1993; Utter and Allendorf 1994; Kitano et al. 1997; Wilson and Turner 2009), which are unable to reliably identify introgression or resolve historically “difficult” nodes. Genetic loci sampled across the whole genome provide sufficient sample size to distinguish gene-tree species-tree discordance caused by incomplete lineage sorting (ILS) vs. introgression, enabling evolutionary histories to be represented as networks (which include reticulate edges in addition to bifurcations, representing both vertical descent and gene flow between lineages) rather than bifurcating trees (Edelman et al. 2019; Suvorov et al. 2022a; Tricou et al. 2022; Ji et al. 2025). Currently, seven Pacific salmon genomes (*O. keta*, *O. nerka*, *O. kisutch*, *O. tshawytscha*, *O. gorbuscha*, *O. masou masou*, and *O. masou ishikawae*), and four trout genomes are available (*O. mykiss*, *O. henshawi*, *O. gilae*, and *O. lewisi*; representing the major lineages in the clade). With these genomes we can finally tackle long-standing hypotheses of introgression in *Oncorhynchus* (Fig. 1C).

To determine how introgression has shaped the evolutionary history of *Oncorhynchus*, we leveraged available chromosome-level genome assemblies to estimate the dated species tree, infer a species network, conduct ancestral state reconstruction of historical ranges to determine how lineages would have overlapped at the time of gene flow, and investigate putative genetic barriers to introgression, specifically mito-nuclear incompatibilities and post-zygotic structural barriers in the genome such as chromosomal inversions. This work suggests that deep ancestral introgression has occurred in *Oncorhynchus*, but mito-nuclear incompatibilities, and the resulting decrease in hybrid fitness, have possibly prevented the *O. gorbuscha* + *O. keta* + *O. nerka* clade from introgressing with other lineages in *Oncorhynchus*.

Furthermore, we do not find evidence of introgression between sampled rainbow and cutthroat trout lineages that predates human hatchery practices. These findings have important implications for the management of this economically and culturally vital lineage.

## Materials and Methods

### Dataset

We sampled from 11 publicly available genomes of *Oncorhynchus*. This included, from the Pacific trout, the westslope cutthroat trout (*O. lewisi*; Flores et al. 2025; which was recently elevated to species level from *O. clarkii lewisi* [Page et al. 2023]), the Lahontan cutthroat trout (*O. henshawi*; Payne et al. 2025), the Columbia River redband trout/rainbow trout (*O. mykiss gairdneri*; US Department of Agriculture 2023) and the Gila trout (*O. gilae*; Barela Hudgell et al. 2026). And from the Pacific salmon, our sampling included the Chinook salmon (*O. tshawytscha*; Christensen et al. 2018), the chum salmon (*O. keta*; Rondeau et al. 2023a), the coho salmon (*O. kisutch*; Rondeau et al. 2023b), the sockeye salmon (*O. nerka*; Christensen et al. 2020), the pink salmon (*O. gorbuscha*; Christensen et al. 2021), the masu/cherry salmon (*O. masou masou*; Christensen et al. 2025), and the amago salmon (*O. masou ishikawae*; Center for Ecological Research, Kyoto University 2023). The two subspecies of masu salmon (*O. masou*) were included as they do not share the same sex chromosome, the consequences of which merit further study (Christensen et al. 2025). We used the genome assembly of *Salvelinus alpinus* (Kurta et al. 2025) as an outgroup. We note the genome assemblies of *O. tshawytscha* (Christensen et al. 2018), *O. mykiss gairdneri* (US Department of Agriculture 2023), and *O. kisutch* (Rondeau et al. 2023b) were generated from inbred homozygous individuals.

By restricting our dataset to only whole genome assemblies, we omit the Pacific trout *O. apache*, *O. chrysogaster*, *O. aguabonita*, and *O. virginalis*, and the salmon *O. biwaensis* and *O. kawamurae*, for which no genomes have been sequenced. So-called “ghost” introgression from unsampled lineages (due to either a lack of data and/or extinction) to sampled lineages can cause the incorrect inference of introgression between sampled species (Tricou et al. 2022). Our taxon sampling here minimizes this risk. The species *O. mykiss*, *O. aguabonita* (unsampled), *O. apache* (unsampled), and *O. gilae* are all closely related and may be more appropriately viewed together as one species complex (Crête-Lafrenière et al. 2012; Campbell et al. 2023); *O. virginalis* is sister to both *O. lewisi* and *O. henshawi* in the broader cutthroat trout complex (Loxterman and Keeley 2012), while *O. kawamurae* is closely related to *O. nerka* (Muto et al. 2013), and *O. biwaensis* to *O. masou* (Fujioka et al. 2025). Gene flow from these unsampled species to any of our sampled species is expected to manifest as gene flow from the sister of the unsampled species, although we are unable to detect gene flow to our unsampled species.

### Divergence time estimation and estimation of nuclear substitution rates

#### Ortholog inference

To infer homologous sequences sequences for species tree, divergence time estimation, and species network estimation, we first aligned the 12 genomes (*O. lewisi*, *O. henshawi*, *O. mykiss*, *O. gilae*, *O. tshawytscha*, *O. keta*, *O. nerka*, *O. kisutch*, *O. gorbuscha, O. masou masou*, *O. masou ishikawae*, and the outgroup *S. alpinus*) using Progressive Cactus v2.9.9 (Armstrong et al. 2020) using default alignment settings. We converted the resulting hal file to maf format with hal2maf v2.1 from HALtools (Hickey et al. 2013), with the flag `--maxBlockLen 10000`. We then extracted all loci between 1,000 and 2,000 bp from the maf alignment. We chose this range to balance the two competing requirements of coalescent-based inference (Mirarab 2019). Loci must be long enough to contain sufficient variable sites to estimate a well-resolved gene tree, since gene-tree estimation error is a major source of error for summary species-tree methods and inflates apparent gene-tree–species-tree discordance, the signal on which our introgression tests depend. Conversely, the multispecies coalescent model (Rannala and Yang 2003; Degnan and Rosenberg 2009) assumes no recombination within a locus and free recombination between loci (Flouri et al. 2020), an assumption that degrades as loci lengthen. Simulations indicate that inference of species trees, divergence times, and introgression parameters under the multispecies coalescent is robust to realistic rates of intralocus recombination (Zhu et al. 2022); we therefore treated 2,000 bp as a conservative upper bound. We retained only loci in which gap characters comprised no more than 20% of the alignment, averaged across all taxa.

#### Species tree inference

For each extracted locus we estimated the maximum likelihood gene tree with IQ-TREE v3.0.1 (Wong et al. 2025), using the flags -B 1000 and -m MFP to conduct 1000 ultrafast bootstrap replicates per gene tree (Hoang et al. 2018), and test for the best fitting nucleotide substitution model for each locus (Kalyaanamoorthy et al. 2017). From the resulting bootstrap-support-annotated maximum likelihood gene trees, we estimated the species tree with weighted ASTRAL v1.23.3.7 (Zhang and Mirarab 2022), which weights quartets by gene tree branch support, specifying *S. alpinus* as the outgroup. Branch support on the species tree was quantified as local posterior probability (Sayyari and Mirarab 2016).

#### Divergence time estimation

Because variation in substitution rate among lineages can produce false-positive signals of introgression (Frankel and Ané 2023), and because divergence times are needed to evaluate whether hybridization between lineages was temporally feasible, we estimated divergence times and lineage-specific substitution rates in BEAST v2.7.7 (Bouckaert et al. 2014, 2019). We concatenated the 100 species-tree-concordant loci with the highest average bootstrap support (> 65%) and analyzed them under GTR+Γ☐(Tavaré 1986; Yang 1994) with an optimized relaxed clock (Drummond et al. 2006; Zhang and Drummond 2020) and a birth–death tree prior (Gernhard 2008; Douglas et al. 2021). Following Lecaudey et al. (2018), we applied uniform calibrations with a maximum root age of 25 Ma and fossil-based minimum ages of 11 Ma for crown Pacific salmon, 6 Ma for *O. gorbuscha* + *O. keta* + *O. nerka*, and 3.2 Ma for Pacific trout (Eiting and Smith 2007; Crête-Lafrenière et al. 2012). Four independent runs were combined after burn-in and summarized as a maximum clade credibility tree (See Supplementary Note S1 for further details and justifications of the divergence time analysis).

To further validate recovered divergence times, we utilized the Pairwise Sequential Markovian Coalescent (PSMC) in lineages expected to have shared a Most Recent Common Ancestor (MRCA) within the past 5 million years, and are thus expected to coalesce within this time frame (see Supplementary Note S2 for implementation of the PSMC).

### Species network inference

Approaches to detecting introgression trade off scalability, model assumptions, and the forms of introgression they can detect, so we combined complementary methods. GhostParser (Tolman and Suvorov 2025) and the Discordant Attachment Frequency Tool (DAFT) (Mishra et al. 2026) scale to genome-wide sets of gene trees and localize introgression to specific branches of the species tree, with GhostParser also distinguishing ghost from sampled introgression, while MPL in PhyloNet (Yu and Nakhleh 2015) infers an explicit network with inheritance probabilities. All three, however, rely on estimated gene trees. We therefore tested candidate events under the full-likelihood MSC-M model in BPP (Flouri et al. 2023), which accounts for gene-tree uncertainty but is computationally demanding and requires a priori hypotheses. Finally, we applied the PSMC to the genome of an F1 *O. mykiss* × *O. lewisi* hybrid, which detects gene flow as parental coalescence predating speciation, independent of gene-tree discordance.

#### Mapping introgression events onto the species tree

We used GhostParser v1.0 and DAFT v1 to map introgression events onto our estimated species tree. GhostParser applies the Discordant Counts Test (DCT; Suvorov et al. 2022a,b) to gene-tree triplets and compares the heights of concordant and discordant topologies to identify the recipient of introgression and whether the donor is a sampled or ghost lineage. We used the coalescent species tree as the guide tree, the bootstrapped gene trees as input, and *S. alpinus* as the outgroup (Supplementary Note S3 for further detail).

DAFT counts pairs of species-tree branches that appear as sisters in gene trees (discordant attachments) and tests, for each branch, whether another lineage attaches to it more frequently in gene trees than would be expected under ILS alone (Mishra et al. 2026). Using DAFT, we implemented both the sister-branch test and the avuncular (“uncle”) test with short-branch/non-coalescence parameters enabled and inferred the direction of gene flow with djiNNI. We considered attachments significant when the z-score of the corrected uncle test was below −1.96 (P < 0.05), or, for pairs where the uncle test could not be applied, that of the corrected sister test (Mishra et al. 2026). We inferred species networks with InferNetwork_MPL (MPL; Yu and Nakhleh 2015; Cao and Nakhleh 2019) in PhyloNet v0.3.0 (Than et al. 2008) from all estimated gene trees, allowing zero to four reticulations with four replicate runs each, and selected the best network by the Akaike Information Criterion (AIC; Akaike 1974) among the top networks from each run (Supplementary Note S3).

#### Gene Flow Hypothesis Testing

To further test the putative introgression events identified with GhostParser, DAFT, and MPL, we fit four Multispecies-Coalescent-with-Migration (MSC-M) models in BPP v4.7.0 (Flouri et al. 2020, 2023) to our locus alignments. Models 1–3 tested candidate events recovered by these methods, and model 4 explicitly tested gene flow between rainbow and cutthroat trout lineages, given its conservation importance (Supplementary Note S4 for full model details). Following Ji et al. (2025), we assigned migration rates the prior M ∼ G(2, 10), ran each model twice, and assessed convergence with ESS > 200 and R☐< 1.1. We evaluated each migration parameter with Bayes factors (B☐☐) calculated with the Savage–Dickey ratio (Ji et al. 2023, 2025) at ε = 0.001 and ε = 0.0001, retaining events with B☐☐> 100 (Kass and Raftery 1995).

#### Further introgression testing using the genome of an F1 hybrid

The publicly available genome assembly of an F1 hybrid of *O. mykiss irideus* and *O. lewisi* (Page et al. 2023) offers another avenue to test for introgression between these lineages (see Supplementary Note S5 for theoretical rationale).

To implement the PSMC on this F1 hybrid, we mapped the ONT reads from this sequencing project (Page et al. 2023) to the reference genome assembly of *O. henshawi* with minimap2 v2.1 (Li 2018), using the flag `map-ont`. We then conducted variant calling and implemented the PSMC as described in Supplementary Note S2.

### Putative barriers to introgression

#### Ancestral state range reconstruction

To model whether putatively introgressing clades would have shared ranges at the time of hypothesized introgression, and to determine if the lack of ancestral introgression in other clades of *Oncorhynchus* could have been caused by historically disjunct ancestral ranges, we conducted ancestral range estimation with BioGeoBEARS v1.1.3 (Matzke 2013, 2014). We used our time-calibrated phylogeny as the input tree, and defined five broad geographic areas, spanning the range of sampled *Oncorhynchus*: Japan, Northeast Asia, Western North America (considered to be west of the Cascade and Sierra Nevada mountain ranges), Intermountain West (considered to be east of the Cascade and Sierra Nevada mountain ranges and north of the Mojave Desert), and Southwest North America (considered to be east of the Cascade and Sierra Nevada mountain ranges and south of the Mojave Desert). We only allowed for distribution through contiguous states, considering Japan as adjacent to Northeast Asia, Northeast Asia as adjacent to Western North America, Western North America as adjacent to both the Intermountain West and Southwest North America, but the Intermountain West and Southwest North America as not adjacent to each other. We fit the Dispersal-Extinction-Cladogenesis (Ree and Smith 2008), Dispersal-Vicariance-like (Ronquist 1997; Matzke 2014), and BayArea-like (Landis et al. 2013; Matzke 2014) models, all with and without the “Jump” speciation parameter J (Matzke 2014) for a total of six models, which we compared using AIC, Akaike Information Criterion corrected (AICc), and Bayesian Information Criterion (BIC) weights (Matzke 2022).

### Mitochondrial-nuclear genome incompatibility

Mitochondrial-nuclear genome incompatibility (MNI) is another potential barrier to introgression. Inter-specific hybrids inherit mitochondrial and nuclear genomes from two different lineages. If the genes from the nuclear and mitochondrial genomes are incompatible and cannot properly support oxidative phosphorylation, hybrids will have decreased fitness, or may be inviable, posing a major barrier to introgression between species (Burton et al. 2013).

To investigate MNI in *Oncorhynchus*, we compared the mitochondrial genomes of all our sampled species except *O. lewisi*, for which a mitochondrial genome assembly is not available. We retrieved the mitochondrial genome sequences for these taxa from GenBank (Table S1) and aligned the sequences using MUSCLE with default parameters (Edgar 2004) in AliView (Larsson 2014). Because mitochondrial genomes are circular and the GenBank sequences did not share a common start position, we then manually rotated sequences whose ends were offset so that all began at the same position. We estimated a maximum likelihood tree from this alignment with IQ-TREE v3.1.1 (Wong et al. 2025), utilizing the flags `--b 1000` and `-m TEST` to conduct 1000 bootstrap replicates from the alignment, and test for the best-fitting nucleotide substitution model.

We primarily considered mitochondrial divergence among three clades: (i) *O. keta*, *O. gorbuscha*, and *O. nerka*; (ii) *O. kisutch* and *O. tshawytscha*; and (iii) Pacific trout (*O. mykiss*, *O. gilae*, *O. henshawi*). Groups ii and iii represent the lineages inferred to have introgressed, whereas group i showed no evidence of introgression with any other lineage.

To estimate lineage-specific substitution rates in the mitochondrial genome, we dated the mitochondrial genome alignment with MCMCtree in PAML v4.10.10 (Yang 2007) on two fixed topologies rooted with *S. alpinus*: the BEAST species tree, pruned to the taxa in the mitochondrial dataset, and the maximum-likelihood mitochondrial genome tree. We used the approximate likelihood method (dos Reis and Yang 2011) under GTR+Γ (Tavaré 1986; Yang 1994) and an independent log-normal relaxed clock (Rannala and Yang 2007), with soft-bounded calibrations (Yang and Rannala 2006) set to the 95% highest posterior density (HPD) intervals of node ages in the BEAST species tree. On the mitochondrial topology, the most recent common ancestor of Pacific salmon and the node uniting *O. kisutch* + *O. tshawytscha* with masu salmon were left uncalibrated. Estimates for each topology were obtained from two independent chains combined after burn-in (Supplementary Note S6).

To evaluate selective constraint, we aligned the 13 mitochondrial protein-coding genes in a codon-aware manner (3,801 codons; Supplementary Note S6) and estimated branch-specific ratios of nonsynonymous to synonymous substitution rates (dN/dS, ω) on both topologies with the free-ratio model (Yang 1998) in codeml in PAML v4.10.10, under the codon substitution model of Goldman and Yang (1994) with F3x4 codon frequencies. We tested for among-branch variation in ω by comparing the free-ratio model with the one-ratio model (M0; a single ω across the tree) using likelihood-ratio tests (LRTs; df = 18) for the concatenated alignment and each gene, with Holm (1979) correction across genes. Each model was run from at least two starting values until two runs converged on the same maximum likelihood (Supplementary Note S6).

To visualize divergence across individual gene features, we calculated the mean uncorrected pairwise distance (*p*-distance; Nei and Kumar 2000) across all between-group sequence pairs in 251-column sliding windows, treating gaps as mismatches. To identify regions in which groups ii and iii were more similar to each other than either was to group i, we calculated for each window Δ = ½[*d*(i,ii) + *d*(i,iii)] − *d*(ii,iii), where positive values indicate that groups ii and iii are closer to each other than to group i. Runs of ≥ 80 columns with Δ ≥ 0.02 were retained as candidate regions and mapped onto gene features. To assess whether these regions reflected elevated divergence within a single group, we also calculated, for each group, the mean proportion of sequences differing from the majority-rule consensus of the ten *Oncorhynchus* sequences in each window. Calculations were conducted with NumPy v2.2 (Harris et al. 2020).

#### Rearrangements in genome architecture

To identify inversions, transpositions, or duplications that could have presented post-zygotic barriers between the major lineages (Pacific trout; masu salmon; *O. kisutch* and *O. tshawytscha*; *O. keta*, *O. gorbuscha*, *O. nerka*), we used halSummarizeMutations v2.1 from HALtools (Hickey et al. 2013) to identify these rearrangements in ancestral genomes of each of our four focal lineages, and the internal branch leading to the ancestor of Pacific salmon.

To test for historical chromosomal fissions and fusions that could have created post-zygotic barriers, we modeled chromosome-number evolution with ChromEvol under six candidate rate models spanning combinations of ascending dysploidy, descending dysploidy, duplication, demi-duplication, and base-number transitions, implemented with the default model parameters (Glick and Mayrose 2014; Rice and Mayrose 2021). Model support was compared by AICc.

## Results

### Divergence time estimation

#### Ortholog inference and coalescent species tree estimation

We recovered 2,486 alignment blocks within our size range, in which each of the 12 taxa was represented by exactly one sequence. All relationships in the coalescent (ASTRAL) species tree had strong posterior support (posterior probability = 1), including the monophyletic Pacific trout and Pacific salmon clades (Fig. S1). Within Pacific trout, the rainbow and cutthroat species complexes were recovered as expected: *O. mykiss* was sister to *O. gilae*, and *O. lewisi* was sister to *O. henshawi*. Within Pacific salmon, the two masu salmon subspecies were sister to all other species, which formed two clades: *O. kisutch* + *O. tshawytscha*, and *O. nerka* + (*O. gorbuscha* + *O. keta*).

#### Divergence time estimation and nucleotide substitution rate inference

Our divergence time estimates between *Salvelinus* and *Oncorhynchus*, Pacific trout and Pacific salmon, and masu and all other Pacific salmon agree well with the previous fossil calibration analyses (Lecaudey et al. 2018; Fig. 2A). Nucleotide substitution rates across the tree are largely consistent (0.0005-0.0009 substitutions/site/Myr), with *O. gorbuscha* being one notable outlier (0.001 substitutions/site/Myr). Under the PSMC, *O. henshawi* and *O. lewisi* do not appear to coalesce by ∼5 Ma (Fig. 2B), consistent with a divergence time falling between 4 and 9 Ma in the BEAST analysis (Fig. 2A).

Coalescence times estimated for Pacific salmon in BEAST do not overlap with the estimated time of coalescence under the PSMC. Under the PSMC the two masu salmon coalesce within 500,000 years but are estimated to have diverged > 2 Ma in BEAST (Fig. 2A,C). The masu salmon then coalesce with *O. nerka* at ∼2 Ma, but are estimated to have diverged > 10 Ma in the dated species tree (Fig. 2A,C).

**Figure 2.**
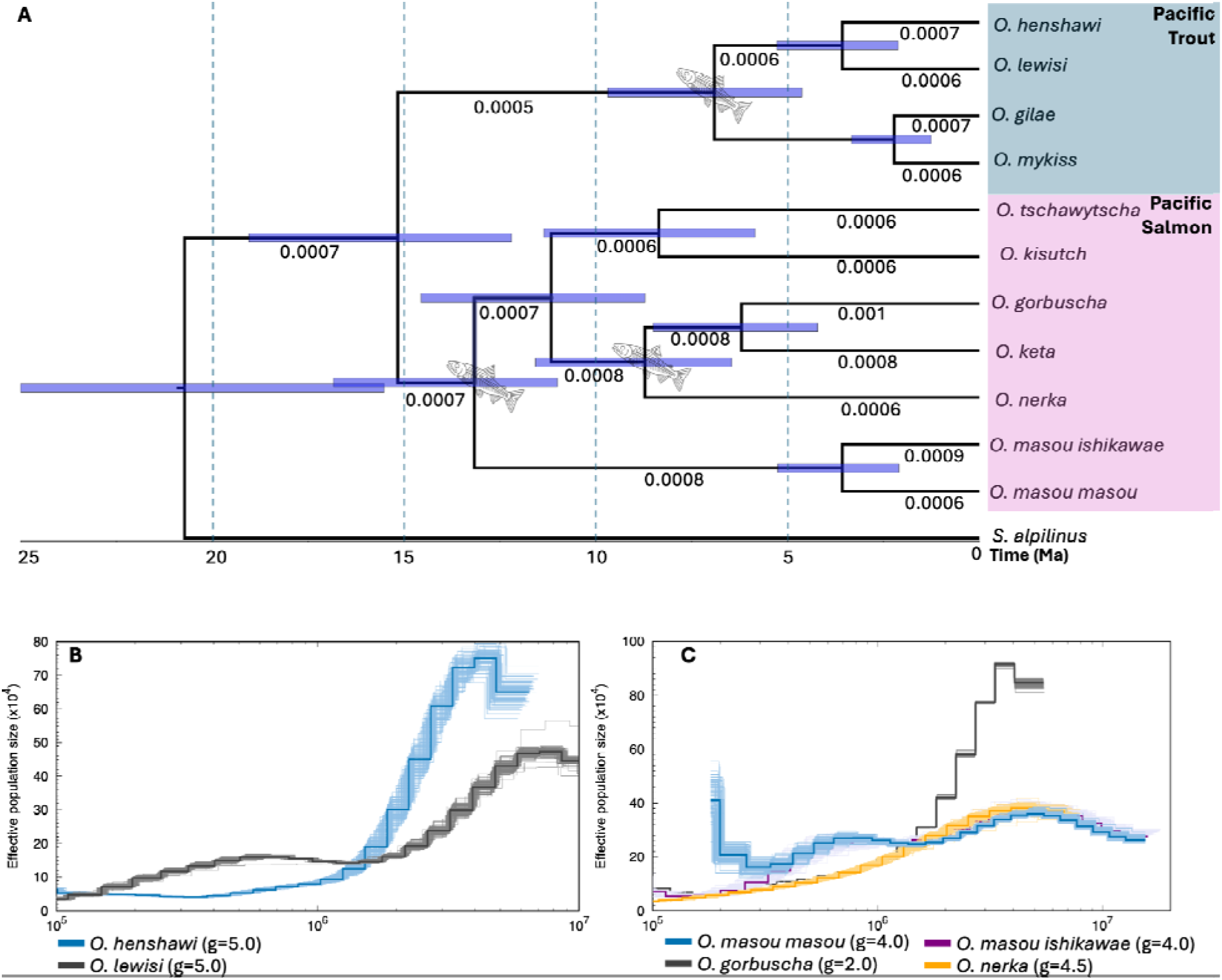
Divergence times and demographic histories of Oncorhynchus. A) Divergence times from the maximum clade credibility tree, with the 95% highest posterior density interval displayed as blue bars at each node. Fossil calibrations on the tree are also shown with a fish symbol. O. rastrosus, O. ketopsis, and O. lacustris were used to set the minimum age constraints for the Pacific salmon; O. gorbuscha, O. keta, and O. nerka; and Pacific trout. Branches are annotated with the recovered nucleotide substitution rates. B) Demographic histories of O. henshawi (blue) and O. lewisi (grey), modeled with the Pairwise Sequential Markovian Coalescent (PSMC). C) Demographic histories of O. masou masou (blue), O. masou ishikawae (purple), O. gorbuscha (grey), and O. nerka (orange), modeled with the PSMC. O. masou ishikawae rapidly coalesces with O. masou masou, and falls under this model for much of the plot. Note a genome-wide mutation rate of 3e-9 was used for each species, with generation times shown in the legend. The coalescence times from PSMC models are identified by points of shared overlap in demographic histories.

### Historical introgression

**Figure 3.**
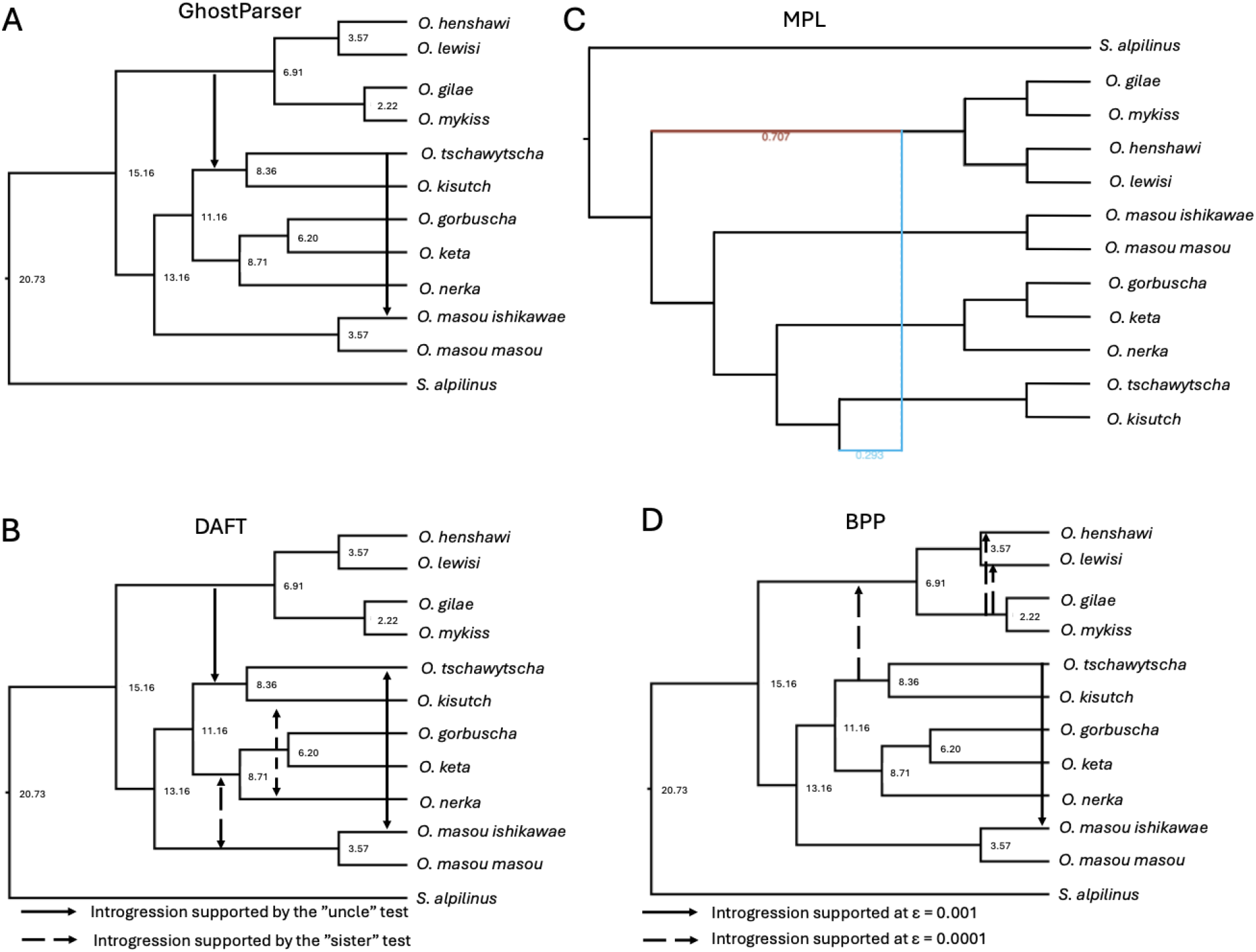
Historical introgression within Oncorhynchus. Introgression events identified by (A) GhostParser (solid black lines represent gene flow identified with GhostParser) and (B) DAFT projected onto the dated species tree. Although the tree is dated, the introgression events are not. (C) The species network recovered with MPL from PhyloNet, with branch lengths converted into millions of years. The inheritance probabilities for each of the two divergent network paths are shown. The brown line is representative of the “non-introgressed” pathway, while the blue line represents the introgressed pathway, jumping into the ancestor of O. tshawytscha and O. kisutch as a result of gene flow from the ancestor of O. kisutch and O. tshawytscha to the ancestor of O. gilae and O. lewisi. (D) Migration parameters retained after fitting four separate migration models to the dataset in BPP. Migration parameters not strongly supported after calculating Bayes Factors with Savage-Dickey ratios were dropped.

GhostParser (Fig. 3A; Fig. S2) identified introgression from the ancestral trout lineage to the ancestor of *O. tshawytscha* and *O. kisutch*, and from *O. tshawytscha* to *O. masou ishikawae*. It also identified *O. masou masou* as a putative ghost recipient; however, the alternative hypothesis of *O. masou ishikawae* as the recipient of introgression from *O. tshawytscha* had much higher bootstrap support across loci (Fig. S2).

DAFT (Table S2) estimated eleven total introgression pairs but only four of these pairs had significant support (z-score < -1.96; Fig. 3B). Of the introgression that could be tested with DAFT’s more robust “uncle” test (> 1 nearest-neighbor interchange [NNI] apart), only gene flow from Pacific trout to the ancestor of *O. kisutch* and *O. tshawytscha*, and between *O. tshawytscha* and *O. masou ishikawae*, was retained. Possible introgression identified with the “sister” test (1 NNI apart) included bidirectional introgression between the ancestor of masu salmon and the ancestor of *O. keta*, *O. gorbuscha*, and *O. nerka*; *O. kisutch* and *O. nerka*; and *O. gorbuscha* and *O. nerka*.

The best-fitting MPL model from PhyloNet contained one introgression event, from the ancestor of trout, again to the ancestor of *O. tshawytscha* and *O. kisutch* (with an introgression probability of 0.29) and did not include any instances of “ghost” introgression in the network (Fig. 3C). All parameters from the BPP models showed evidence of convergence (ESS > 200 and R < 1.1 across both replicate runs; Table S3). Only migration from *O. tshawytscha* to *O. masou ishikawae* was retained at ε = 0.001 in our Savage–Dickey ratio tests of the BPP models (Fig. 3D). However, at ε = 0.0001, migration from the ancestor of *O. kisutch* and *O. tshawytscha* to the ancestor of Pacific trout, and from the ancestor of *O. mykiss* and *O. gilae* to both *O. lewisi* and *O. henshawi*, were also retained (Fig. 3D).

We did not identify any change in coalescence rate between *O. mykiss irideus* and *O. lewisi* in applying the PSMC to the F1 hybrid genome, further suggesting that no ancestral introgression has occurred between *O. mykiss* and *O. lewisi*.

### Ancestral state range reconstruction

**Figure 4.**
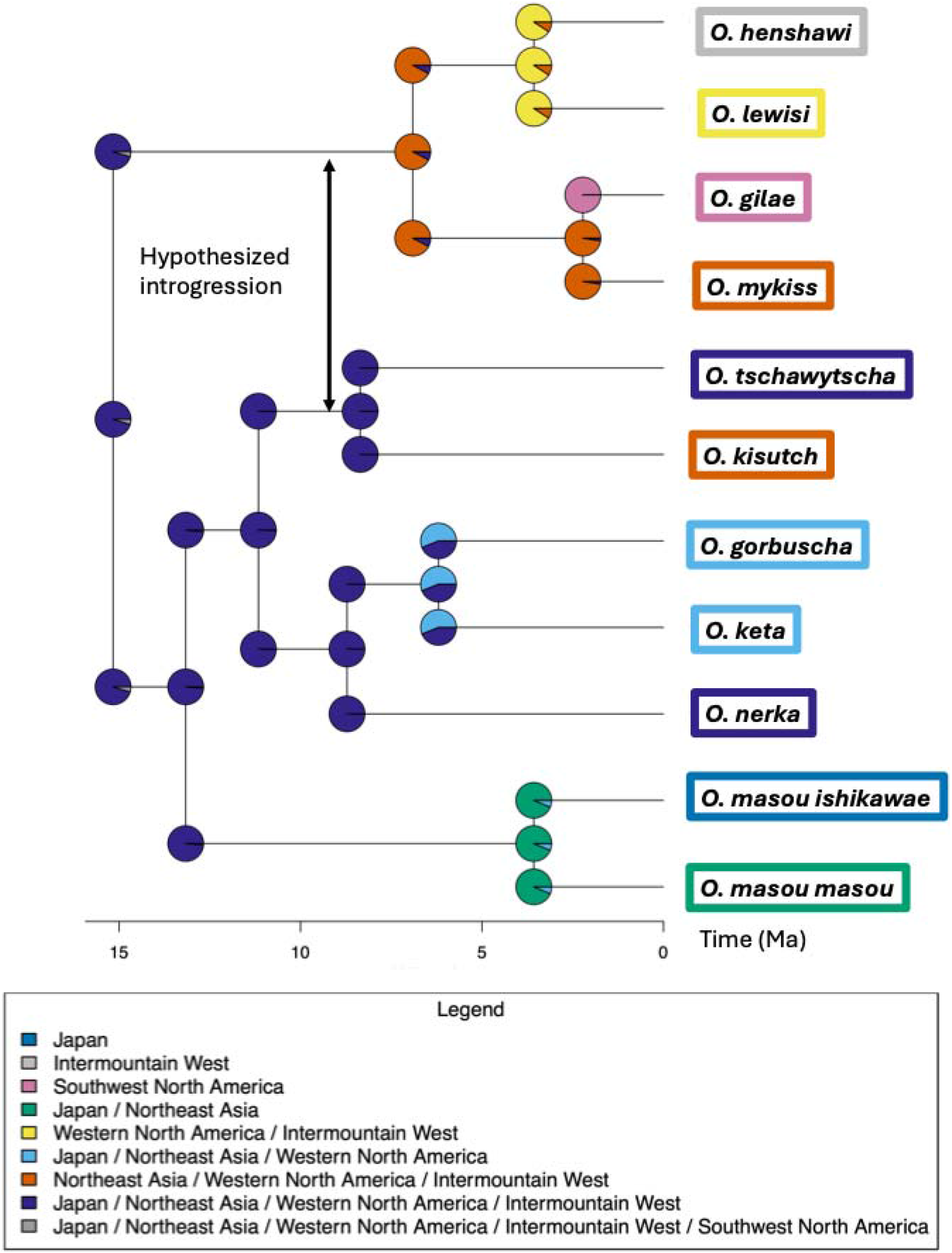
Ancestral range reconstruction of Oncorhynchus. The best-fitting (BAYAREALIKE+J) model is shown with the hypothesized introgression event mapped onto the tree. Tips are colored according to current ranges.

The BAYAREALIKE+J model was strongly supported across all metrics (AICw = 0.93, AICcw = 0.85, BICw = 0.91), with the most probable ancestral range of *Oncorhynchus* spanning Japan, Northeast Asia, Western North America, and the Intermountain West. Under this model, the number of range expansion events per lineage per million years was negligible (d = 1e-7), while each lineage lost an area from its range once every 25 million years on average (e = 0.05). The jump parameter was not weighted especially strongly, with a cladogenesis event probability of founder-event speciation of 0.03. *O. gilae* was the only lineage speciating through a founder event (Fig. 4). The MRCA of *O. tshawytscha* and *O. kisutch* was expected to have retained the ancestral range at the time of putative introgression with the ancestral trout lineage, while the ancestral trout lineage lost Japan between separation from the Pacific salmon and the MRCA of sampled Pacific trout (Fig. 4).

### Genome structure as a potential barrier to introgression

#### Mitochondrial-nuclear genome discordance

While Pacific trout and Pacific salmon were recovered as monophyletic clades in the mitochondrial genome tree, the tree topology did not match the species tree (Fig. S3; Fig. 5A–B). Within the Pacific salmon clade, *O. kisutch* and *O. tshawytscha* were placed as sister to the two subspecies of masu salmon. While the relationships within the three salmon clades were strongly supported in the mitochondrial genome tree, the relationships between them were not (bootstrap support < 90; Fig. S3). *O. kisutch* and *O. tshawytscha* were more similar, by nucleotide composition, to the Pacific trout than to the *O. nerka*, *O. gorbuscha*, and *O. keta* clade, and the Pacific trout mitochondrial genomes were also more similar to masu salmon than they were to *O. nerka*, *O. gorbuscha*, and *O. keta* (Table 1; Fig. S4).

**Table 1.** Mean pairwise mitochondrial nucleotide similarity between clades within *Oncorhynchus*.

|  |  |  |
| --- | --- | --- |
| Clade 1 | Clade 2 | Similarity of mitochondrial genome |
| *O. kisutch* + *O. tshawytscha* | masu salmon | 93.56% |
| Pacific trout | *O. kisutch* + *O. tshawytscha* | 93.38% |
| masu salmon | *O. gorbuscha* + *O. nerka* + *O. keta* | 93.32% |
| Pacific trout | masu salmon | 92.93% |
| Pacific trout | *O. gorbuscha* + *O. nerka* + *O. keta* | 92.3% |
| *O. kisutch* + *O. tshawytscha* | *O. gorbuscha* + *O. nerka* + *O. keta* | 92.18% |
| *NOTE sampled Pacific trout are comprised of *O. henshawi*, *O. mykiss* and *O. gilae*, while sampled masu salmon include *O. masou masou* and *O. masou ishikawae*. |  |  |

**Figure 5.**
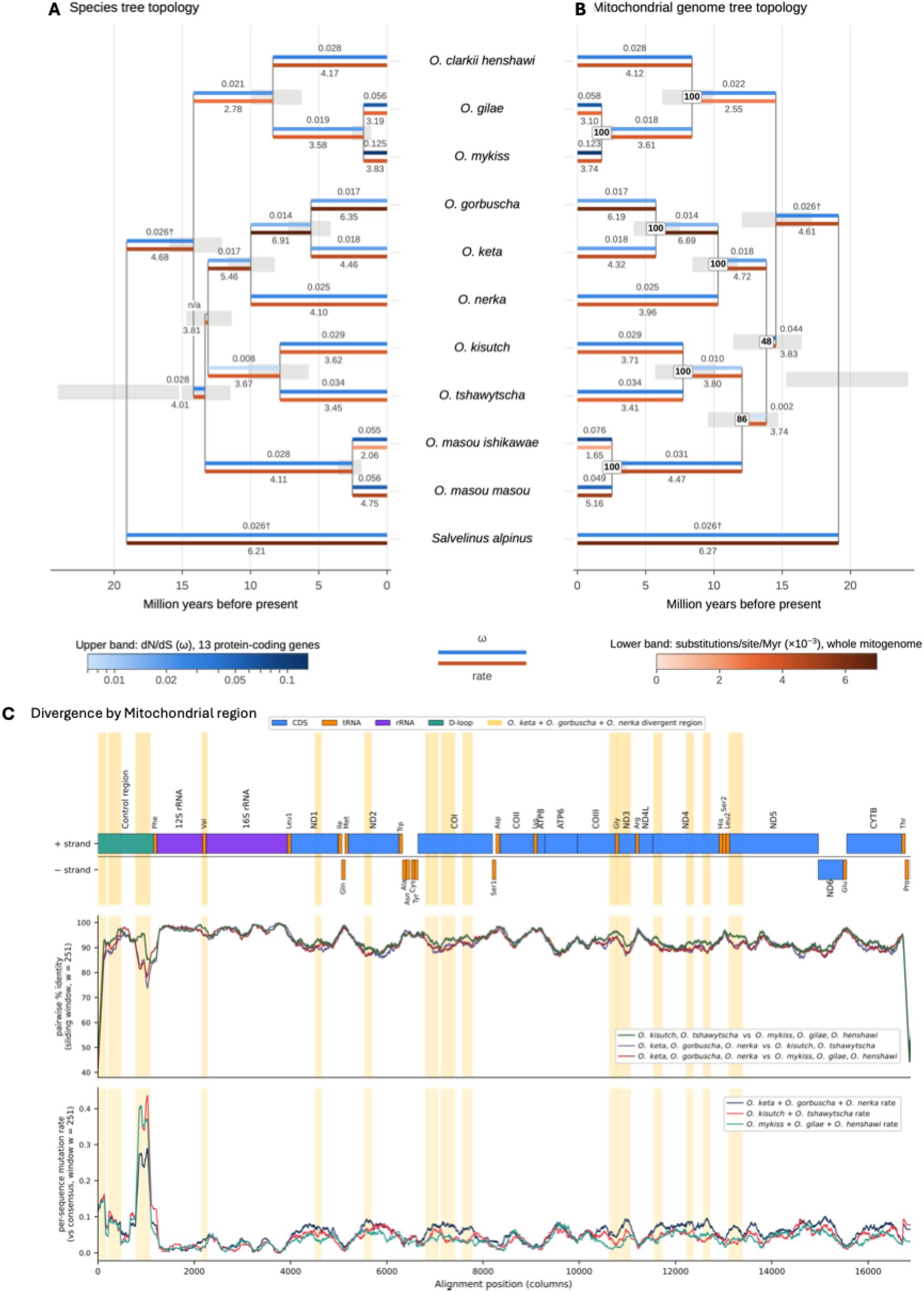
Mitochondrial divergence map between select Pacific salmon and trout. A) Mitochondrial nucleotide substitution rates (estimated with MCMCtree) and dN/dS ratios (ω) per branch assuming the topology inferred from the nuclear genome. B) Mitochondrial nucleotide substitution rates and ω per branch assuming the topology inferred from the mitochondrial genome. In both A) and B) divergence time estimates come from MCMCtree. C) Mitochondrial divergence by region. (Top) Gene-feature track showing protein-coding genes (CDS, blue), transfer RNAs (tRNA, orange), ribosomal RNAs (rRNA, purple), and the control region / D-loop (green) on the plus and minus strands. (Middle) Sliding-window pairwise nucleotide identity (window = 251 columns) between the Pacific trout (O. mykiss + O. gilae + O. henshawi) and O. kisutch + O. tshawytscha (green: the ancestors of these two clades are hypothesized to have introgressed); O. keta + O. gorbuscha + O. nerka and O. kisutch + O. tshawytscha (purple); and O. keta + O. gorbuscha + O. nerka and Pacific trout (red). (Bottom) Per-sequence mutation rate within each group (O. keta + O. gorbuscha + O. nerka [blue]; Pacific trout [green]; O. kisutch + O. tshawytscha [orange]), calculated as the fraction of within-group sequences differing from the column-wise majority base, averaged over the same 251-column window. Beige vertical bars spanning all three panels mark alignment windows in which the O. kisutch + O. tshawytscha and Pacific trout identity exceeds the mean of the two comparisons involving O. keta + O. gorbuscha + O. nerka by ≥ 2.0% for runs of ≥ 80 nucleotides. Sixteen such regions were recovered, concentrated in the control region, the 12S/tRNA-Val/16S boundary, ND1, ND2, COI, COIII–tRNA-Gly–ND3, ND4, tRNA-Leu2, and ND5.

Both MCMCtree analyses converged (ESS > 20,000 and R☐ ≤ 1.0002 for node ages and branch rates). Branch-specific substitution rates were largely consistent between topologies and fell mostly between 3 and 5 × 10⁻³ substitutions/site/Myr (Fig. 5A–B; Table S4). The fastest branches belonged to the stem of *O. gorbuscha* + *O. keta* (6.9 × 10⁻³ [4.2–10.3] on the species topology; 6.7 × 10⁻³ [4.0–10.1] on the mitochondrial topology) and the *O. gorbuscha* terminal branch (6.3 [4.5–8.3]; 6.2 [4.3–8.2]). The slowest branches were the Pacific trout stem (2.8 [1.5–4.3]; 2.6 [1.3–4.0]) and the *O. masou ishikawae* terminal branch (2.1 [1.2–2.9]; 1.7 [0.9–2.4]).

Branch-specific ω estimates indicated strong purifying selection throughout (Fig. 5A–B). On most branches ω was between 0.008 and 0.034, and it was lowest within the *O. keta* + *O. nerka* + *O. gorbuscha* clade (0.014–0.025). ω was elevated on short terminal branches in Pacific trout (O. mykiss 0.125 and 0.123; O. gilae 0.056 and 0.058, species and mitochondrial topologies respectively) and masu salmon (0.049–0.076). Estimates on branches shared by both topologies were nearly identical. The exceptions were branches adjacent to the discordant placement of masu salmon. Allowing ω to vary among branches improved fit over a single ω for the concatenated alignment (M0 ω = 0.026; 2ΔlnL = 61.6 and 59.0 on the species and mitochondrial topologies, *P* < 10☐☐ ; Table S5). Gene-wise M0 ω ranged from 0.003 (ND4L) to 0.050 (ND2), and only ND2 showed significant among-branch variation after correction (Holm-adjusted *P* = 0.044 and 0.007), with ω highest on the *O. mykiss* (0.235 and 0.210) and *O. gilae* (0.152 and 0.168) terminal branches and ≤ 0.039 within the *O. keta* + *O. gorbuscha* + *O. nerka* clade (Table S5). Before correction, ND6 (*P* = 0.040 and 0.022) and CYTB (species topology only; *P* = 0.035) also varied among branches, with the highest ω (excluding branches with ≤ 3 substitutions) on the *O. keta* (0.16 and 0.15) and *O. mykiss* (0.09) terminal branches, respectively (Table S5).

We identified 16 regions in the mitochondrial genome where the percent similarity between *O. kisutch* + *O. tshawytscha* and the Pacific trout exceeds the mean of the similarity of both *O. keta, O. gorbuscha, O. nerka* and Pacific trout, and *O. keta, O. gorbuscha, O. nerka*, and *O. kisutch* and *O. tshawytscha* by ≥ 2.0% for runs of ≥ 80 nucleotides (Fig. 5). These runs were concentrated in the control region, the 12S/tRNA-Val/16S boundary, ND1, ND2, COI, COIII–tRNA-Gly–ND3, ND4, tRNA-Leu2, and ND5 and generally corresponded with regions of an elevated mutation rate in *O. keta, O. gorbuscha,* and *O. nerka* (Fig. 5).

#### Genome structure

We identified seven inversions in both the ancestral Pacific trout and ancestral salmon lineages, both spanning ∼22,000 bp (Table 2). The MRCA of *O. keta*, *O. gorbuscha*, and *O. nerka* was estimated to have undergone only one inversion of 42 bp, while the MRCA of *O. kisutch* and *O. tshawytscha* underwent three inversions totaling 5,880 bp and the ancestral masu salmon had 22 inversions spanning 39,237 bp (Table 2). The MRCA of *O. kisutch* and *O. tshawytscha* also experienced more duplications (∼597,000 duplications totaling 40 million bp against ∼381,000 duplications totaling 23 million bp) and transpositions (186 totaling 254,191 bp against 96 transpositions of 194,300 bp) than the MRCA of *O. keta*, *O. gorbuscha*, and *O. nerka*, although both lineages underwent fewer duplications and transpositions than the MRCA of Pacific trout (∼1.2 million duplications of 72 million bases and 237 transpositions of 307,663 bases) and the MRCA of masu salmon (962,220 duplications of ∼53 million bp, and 155 transpositions of 284,900 bp; Table 2). The ancestral Pacific salmon lineage contained ∼556,000 duplications, totaling ∼26 million bp, and 151 transpositions of 278,182 bp (Table 2).

The best-supported model for chromosome evolution was “Dys”, which models constant single-chromosome ascending and descending dysploidy only (ΔAIC ≥ 3.59 for all alternatives). Under the “Dys” model, per-branch expected rates were 1.14 chromosomal gains and 0.95 chromosomal losses per lineage per million years. The ancestral masu salmon had the highest expected number of fusions and fissions (11.42 gains and 7.95 losses), followed by ancestral trout (7.78 gains and 7.70 losses), the MRCA of *O. kisutch* and *O. tshawytscha* (3.12 gains and 2.34 losses), the MRCA of *O. keta*, *O. gorbuscha*, and *O. nerka* (2.43 gains and 2.22 losses), and the MRCA of Pacific salmon (1.98 gains and 1.76 losses; Table 2).

**Table 2.**
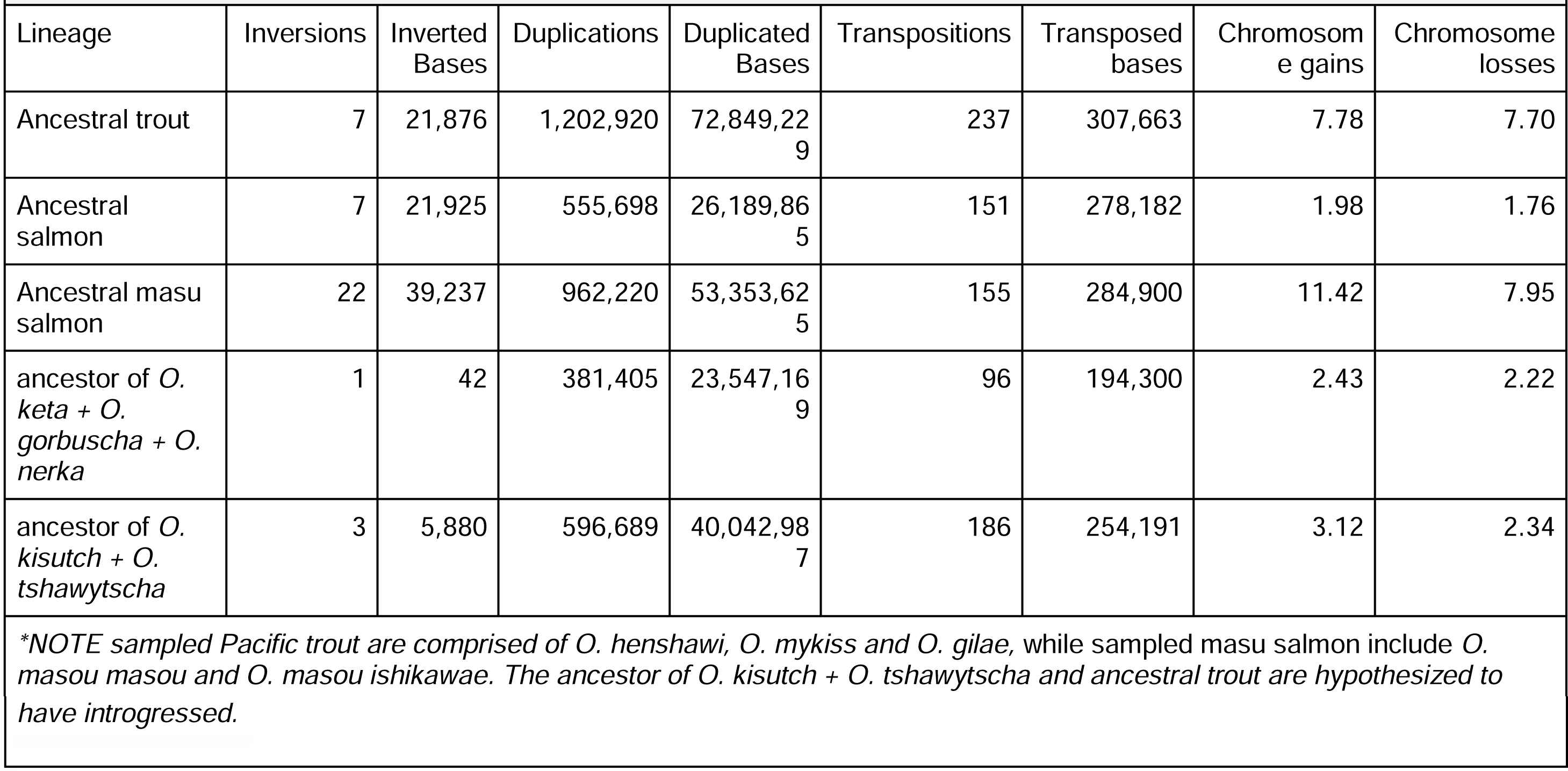
Recovered genome rearrangements within *Oncorhynchus*.

## Discussion

### Deep ancestral introgression between Pacific trout and select Pacific salmon

While the previous divergence time estimates for *Oncorhynchus* from Lecaudey et al. (2018) sampled fewer species, and focused on Salmonidae more generally, our credible intervals for divergence times generally overlap; however, our results tend to skew younger (Fig. 2; e.g., MRCA of *Oncorhynchus* and *Salvelinus*, *Oncorhynchus*, Pacific salmon, and *O. gorbuscha* and *O. keta* are estimated to be between 16–25 Ma, 12–19 Ma, 11–16 Ma, and 4–7 Ma in our study; and 20–23 Ma, 18–22 Ma, 11–13 Ma, and 10.5–12 Ma in Lecaudey et al. 2018). The older ages recovered by Lecaudey et al. (2018) may partly reflect their inclusion of loci discordant with the species tree in a concatenated RADseq dataset, which can inflate branch lengths and thus divergence time estimates (Carruthers et al. 2022). By restricting our dating analysis to loci concordant with the species tree, we expect our estimates to be less affected by this bias. Although they represent, at best, coarse estimates of divergence times, the coalescence times observed in our PSMC analyses fall within the credible intervals of our dated tree (Fig. 2). Thus, these results show that divergence times estimated with the PSMC should be treated with healthy skepticism at present, as fluctuations in genome-wide mutation rates and generation times could both be biasing these estimates.

All four of our tested methods identified introgression between the ancestor of Pacific trout and the ancestor of *O. kisutch* and *O. tshawytscha* (albeit BPP only retained this introgression at ε = 0.0001), although the directionalities differ between methods with GhostParser and DAFT (Table S2) inferring gene flow from trout to the ancestor of *O. kisutch and O. tshawytscha*, but MPL and BPP inferring gene flow in the opposite direction (Fig. 3A–D). As we do not observe elevated nucleotide substitution rates in *O. nerka* and *O. keta* relative to *O. tshawytscha* and *O. kisutch* that could lead to a false introgression signal (Fig. 2; see Frankel and Ané [2023] for a discussion of how rate heterogeneity can lead to incorrect introgression detection; the rate of *O. gorbuscha* is elevated, but GhostParser and DAFT still recover the introgression signal when not sampling this species), and BPP is robust in the presence of such heterogeneity (Frankel and Ané 2023), we did not recover any evidence that this introgression signal is due to nucleotide substitution rate heterogeneity. Furthermore, we do not recover any evidence that this could be an artifact of unsampled “ghost” introgression from either GhostParser or MPL (Fig. 3A,C). These two introgressing lineages also likely inhabited the same, or very similar, geographic regions at the time of introgression (Fig. 4). Thus, the directionality of gene flow may be ambiguous, but the evidence for some sort of introgression between the ancestral trout lineage, and the ancestor of *O. kisutch and O. tshawytscha* is highly compelling, strongly supporting a hypothesis first proposed over thirty years ago (Utter and Allendorf 1994).

DAFT, BPP, and GhostParser identified introgression between *O. tshawytscha* and *O. masou ishikawae* (Fig. 3B; Table S2), and our ancestral state reconstruction showed that *O. tshawytscha* has likely always included Japan in its range, making this introgression reasonably well supported. However, the nucleotide substitution rate of *O. masou ishikawae* is higher than that of *O. masou masou* (Fig. 2), thus we cannot entirely rule out this putative gene flow as an artifact of rate variation. Given the relatively recent divergence between sampled masu salmon, further exploration of this question is more robustly suited to population-level resampling. DAFT also identifies three additional introgression events between lineages too close in the tree for application of the more robust “uncle” test. As these introgression events were not identified in any other tests, we consider them to be unreliable. We do not identify any introgression from the sister of any of our unsampled lineages to another sampled lineage, and can thus conclude that these lineages are not serving as major introgression donors, but we cannot rule out the possibility that they might be introgression recipients at present.

### Historical barriers to introgression in Oncorhynchus

While the ancestral introgression between Pacific trout and the ancestor of *O. kisutch and O. tshawytscha* is notable, the lack of introgression between species which are largely currently sympatric (Crawford and Muir 2008) is equally intriguing, as sympatry can be conducive to introgression (Martin et al. 2013; Waters et al. 2023). Our ancestral range reconstruction analysis found that the ranges for *Oncorhynchus* have historically been highly sympatric, so historical allopatry is not a plausible barrier to introgression, except in the case of Pacific trout and masu salmon (Fig. 4). Despite historical sympatry, reproductive isolation may also be maintained by pre- or postzygotic barriers (e.g., gamete incompatibility or spawning timing). More recent (< 10 Ma) inversions in *Oncorhynchus* serve as barriers to gene flow (Ostberg et al. 2013; Flores et al. 2025), but we recover no evidence of historical rearrangements in genome architecture that could have created barriers between ancestral *Oncorhynchus* lineages, with a similar number of inversions, transpositions, duplications, and chromosome gains and losses in the MRCA of *O. gorbuscha*, *O. keta*, and *O. nerka* as in other, putatively introgressing lineages (Table 2).

Mitochondrial-nuclear incompatibilities do appear to be a possible mechanism preventing introgression. The mitochondrial genomes of *O. kisutch* and *O. tshawytscha* are more similar in nucleotide composition to Pacific trout than they are to *O. nerka*, *O. gorbuscha*, and *O. keta* (Table 1; Fig. S4), suggesting selection may have played a role in the maintenance of mitochondrial genome sequences. This does not appear to be an instance of mitochondrial capture, as the Pacific trout and Pacific salmon clades are recovered as monophyletic in the mitochondrial genome tree (Fig. S3; Fig. 5B). A similar phenomenon has been observed in centrarchid fishes (sunfishes and black basses). The species with faster-evolving mitochondrial genomes tend to produce less-viable hybrids (Bolnick et al. 2008), consistent with the elevated mitochondrial divergence we observed in *O. keta*, *O. gorbuscha*, and *O. nerka* (*Fig. 5*).

The hypothesis of MNI is further supported when investigating the specific regions of the mitochondrial genome where *O. nerka*, *O. gorbuscha*, and *O. keta* are more diverged from the mitochondrial genomes of the Pacific trout and both *O. kisutch* and *O. tshawytscha* (Fig. 5), which occur in genes that have specifically been implicated in mitochondrial-nuclear discordance. In perhaps the most well-studied case in Actinopterygii, mismatched nuclear and mitochondrial genomes can be lethal in hybrids of *Xiphophorus birchmanni* and *X. malinche*, with the mitochondrial genes ND2 and ND6, and the nuclear genes (NDUFS5, NDUFA13) that physically contact Complex I subunits (Moran et al. 2024) specifically implicated. ND2 is notably an area of high divergence between *O. nerka*, *O. gorbuscha*, *O. keta*, and other lineages (Fig. 5).

However, it is important to note that this divergence was largely synonymous: ND2 was the only gene with significant among-branch variation in ω, but ω was lowest within the *O. keta*, *O. gorbuscha*, and *O. nerka* clade and highest in Pacific trout (Table S5), indicating strong purifying selection on ND2 in the non-introgressing clade. Before correction, ND6, also implicated in *Xiphophorus*, varied in ω among branches, with its highest reliable ω on the *O. keta* branch (within the non-introgressing clade; Table S5).

These patterns make mito-nuclear incompatibility a promising avenue for further investigation in *Oncorhynchus* but require extensive transcriptomic data to properly annotate the nuclear genomes for each species, to test whether the diverged mitochondrial genes show signatures of accelerated or coevolutionary selection with their interacting nuclear counterparts.

### Current trends in Oncorhynchus and implications for management

Diverging approximately 6.9 Ma (Fig. 2A), rainbow and cutthroat trout have accumulated significant genetic differences over time yet have retained the ability to successfully hybridize, as rainbow and cutthroat trout Robertsonian chromosomal rearrangements have maintained the same number of diploid chromosome arms (n = 104) when they overlap in temporal and spatial spawning (Flores et al. 2025). Future work with these Pacific trout lineages could uncover more recent introgression within each of these complexes (e.g., Campbell et al. 2023). Such work should offer increased resolution into the histories of the currently ambiguous rainbow and cutthroat species complexes. However, our results here do not provide strong evidence that the rainbow trout and cutthroat trout species complexes have introgressed with each other or with another lineage of *Oncorhynchus* since the divergence of *O. mykiss* and *O. gilae*, despite previous datasets with limited nuclear loci suggesting this may have occurred (Campton and Utter 1985; Allendorf and Leary 1988). No signal is found in traditional tests for introgression (Fig. 3), nor in our implementation of the PSMC with the genome of an F1 hybrid between *O. mykiss* and *O. lewisi*.

Understanding introgression between rainbow and cutthroat lineages is important to effective conservation and management of these two lineages. The decline of trout populations can cause a cascade/ripple effect on the surrounding ecosystem (Baxter et al. 2004; Middleton et al. 2013; Koel et al. 2019). Recent introgression has been extensively demonstrated between these lineages, as initiated by the introduction of nonnative *O. mykiss* to the native ranges of locally endemic cutthroat trout by humans (Hitt et al. 2003; Weigel et al. 2003; Rasmussen et al. 2010; Hohenlohe et al. 2011, 2013; Muhlfeld et al. 2014). Although historical introgression has resulted in a diversified phylogeny for this genus, contact driven by human development and landscape change may present unforeseen consequences. Hybridization between these two populations can lead to several maladaptive consequences including the loss of locally adapted traits, reduced fitness, and decreased resilience to environmental change (Weigel et al. 2003; Flores et al. 2025). The repeated introduction of hybridized or backcrossed lineages in small or geographically restricted populations could decrease local genomic adaptations, which could lead to population decline or extirpation (Todesco et al. 2016). Our results suggest that historical (< 2 Ma) introgression between the cutthroat and rainbow trout species complexes should not be treated as a baseline for management.

Further investigation into the distinction between historical and recent introgression will aid in conserving these taxa, as the presence of ancient gene flow does not necessarily indicate that contemporary hybridization is a natural or beneficial process. As historical introgression occurred within ecological contexts that contrast with modern habitat alteration, hatchery stocking initiatives, and climate change, maintaining genetic integrity within hatcheries is an essential component of preserving adaptive variation for imperiled *Oncorhynchus* lineages. The genetic consequences of hatchery supplementation and translocation programs should be evaluated before introducing low-heterozygosity lineages (i.e., *O. mykiss*) into watersheds containing native cutthroat trout lineages.

## Conclusions

The results presented here provide compelling evidence for ancient introgression between the ancestor of trout and the ancestor of the Pacific salmon *O. kisutch* and *O. tshawytscha*, and possible introgression between *O. tshawytscha* and *O. masou ishikawae*. Other hypothesized ancestral introgression events in *Oncorhynchus* are not strongly supported. Our results highlight the value of genome-wide data for conservation monitoring, as historical or recent introgression is difficult to distinguish using a smaller genetic dataset (i.e., molecular markers). We identify mito-nuclear genome discordance as a putative barrier to introgression between species that have largely been sympatric throughout their evolutionary history, although this hypothesis requires further exploration. Of note is a lack of historical introgression between rainbow and cutthroat trout lineages since the divergence of rainbow and Gila trout, a topic with major conservation and management implications, which has long been debated. Our findings suggest that historical hybridization between these lineages should not be treated as a baseline. Repeated genome-wide sampling across spatial and temporal scales should be integrated in management initiatives to establish hybridization thresholds, genetic compatibility of trout populations before translocation, and identify distinct or rare genotypes that require heightened protections.

## Funding

Student participants were funded through the Youth Climate Action Grant from the City of Boise.

## Acknowledgments

The authors are grateful for support from the board of the Conservation Connection Foundation, especially Kristin Gnojewski and Cindy Busche.

## Data Availability Statement

Locus alignments, gene trees, BEAST input and output files, GhostParser, DAFT, PhyloNet and BioGeoBEARS input and output files, mitochondrial alignments, PAML analysis files, and custom scripts are available on Figshare (https://doi.org/10.6084/m9.figshare.34374180).

## Notes

### Competing Interest Statement

The authors have declared no competing interest.

