## Supplementary materials for "Testing Thirty Years of Hypotheses: Ancient Trout–Salmon Introgression and Limited Pre-Anthropocene Gene Flow Between Rainbow and Cutthroat Trout Shaped Pacific Salmonid Evolution"

***Supplementary figures***

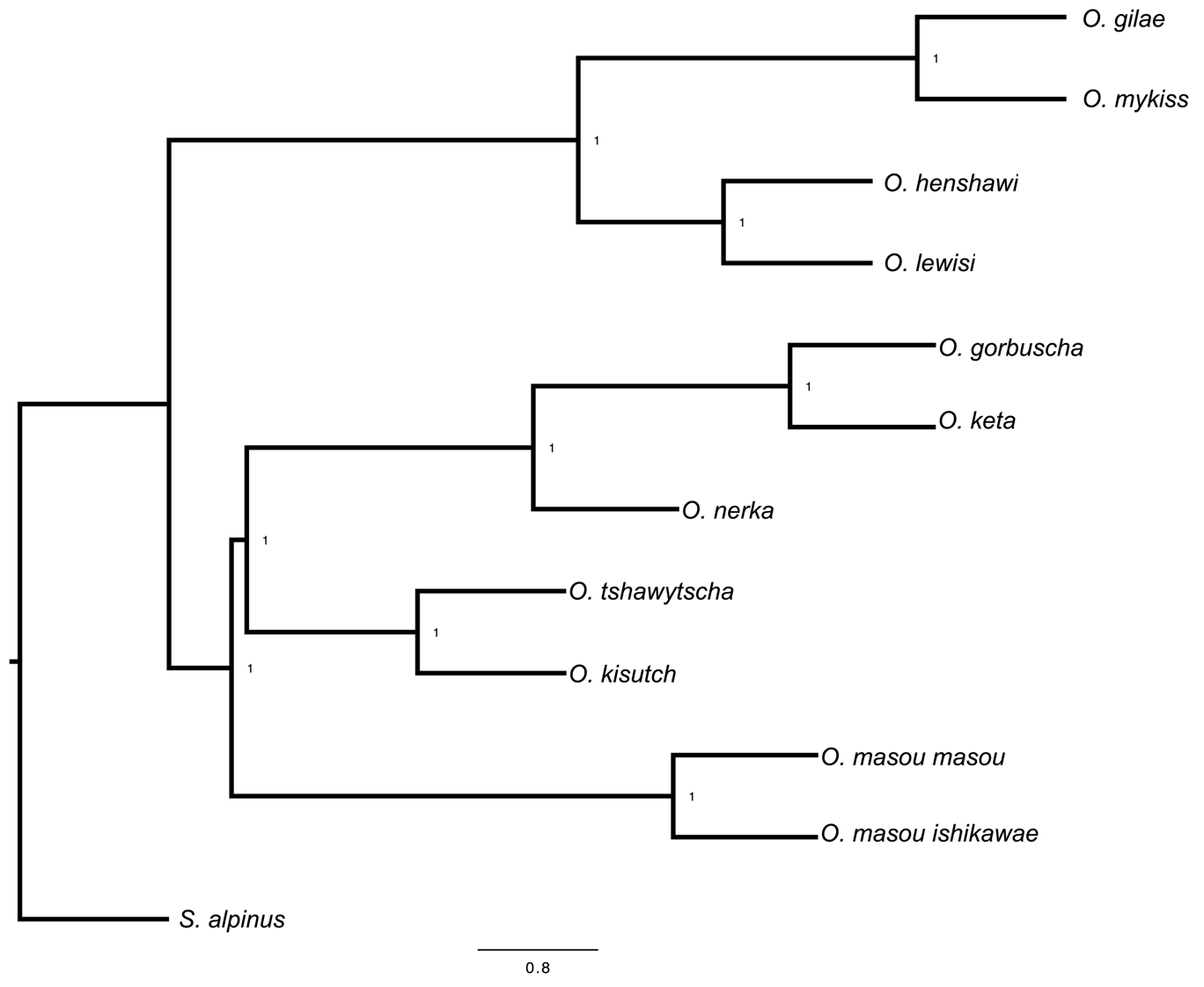

***Supplementary Figure S1. ASTRAL species tree of Oncorhynchus.*** *Posterior probabilities are shown at each node.*

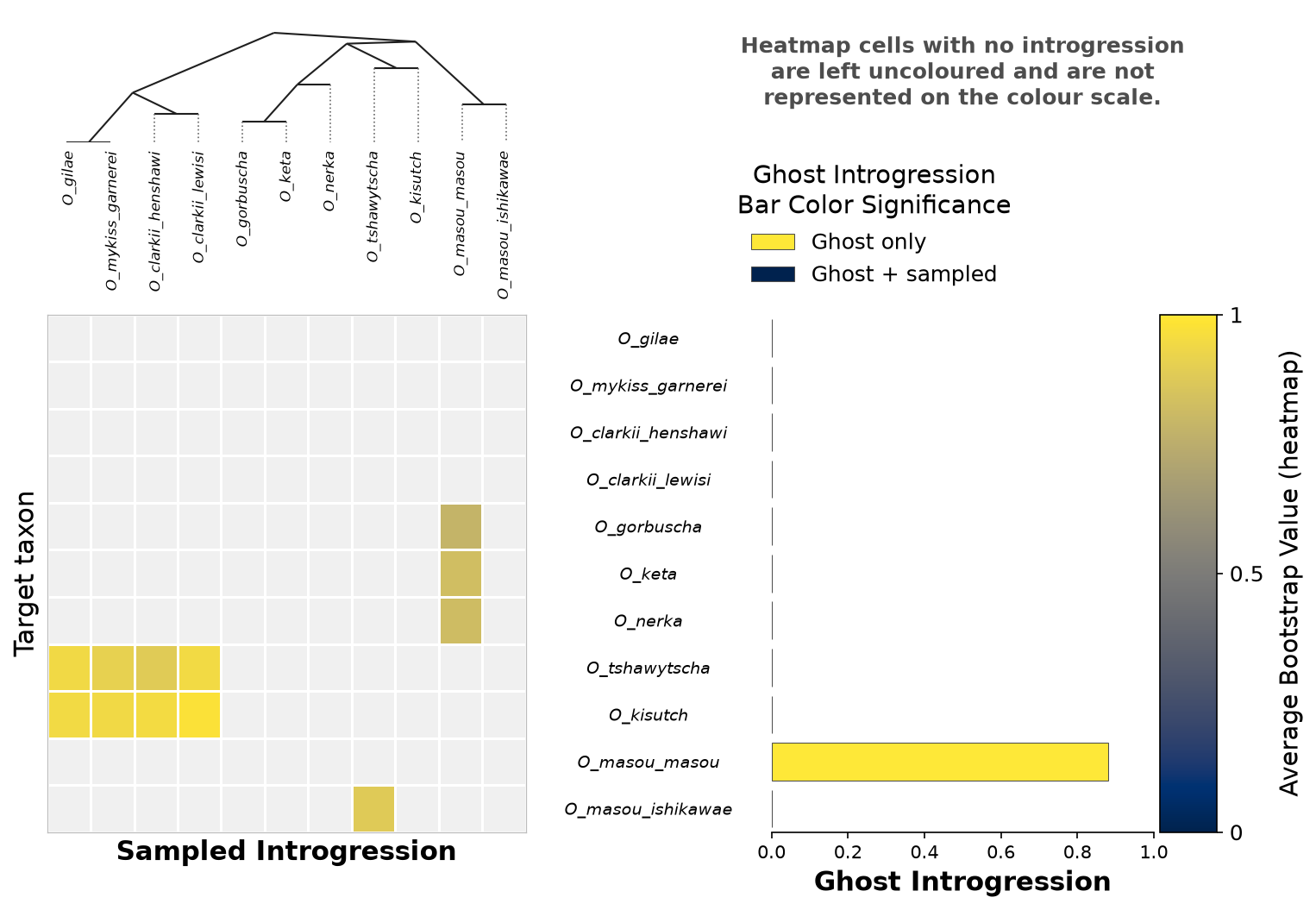

***Supplementary Figure S2. GhostParser support for introgression between sampled and ghost lineages.*** *Bootstrap support is shown for introgression from each donor to target taxon, alongside bootstrap support for possible “ghost” introgression into each extant lineage.*

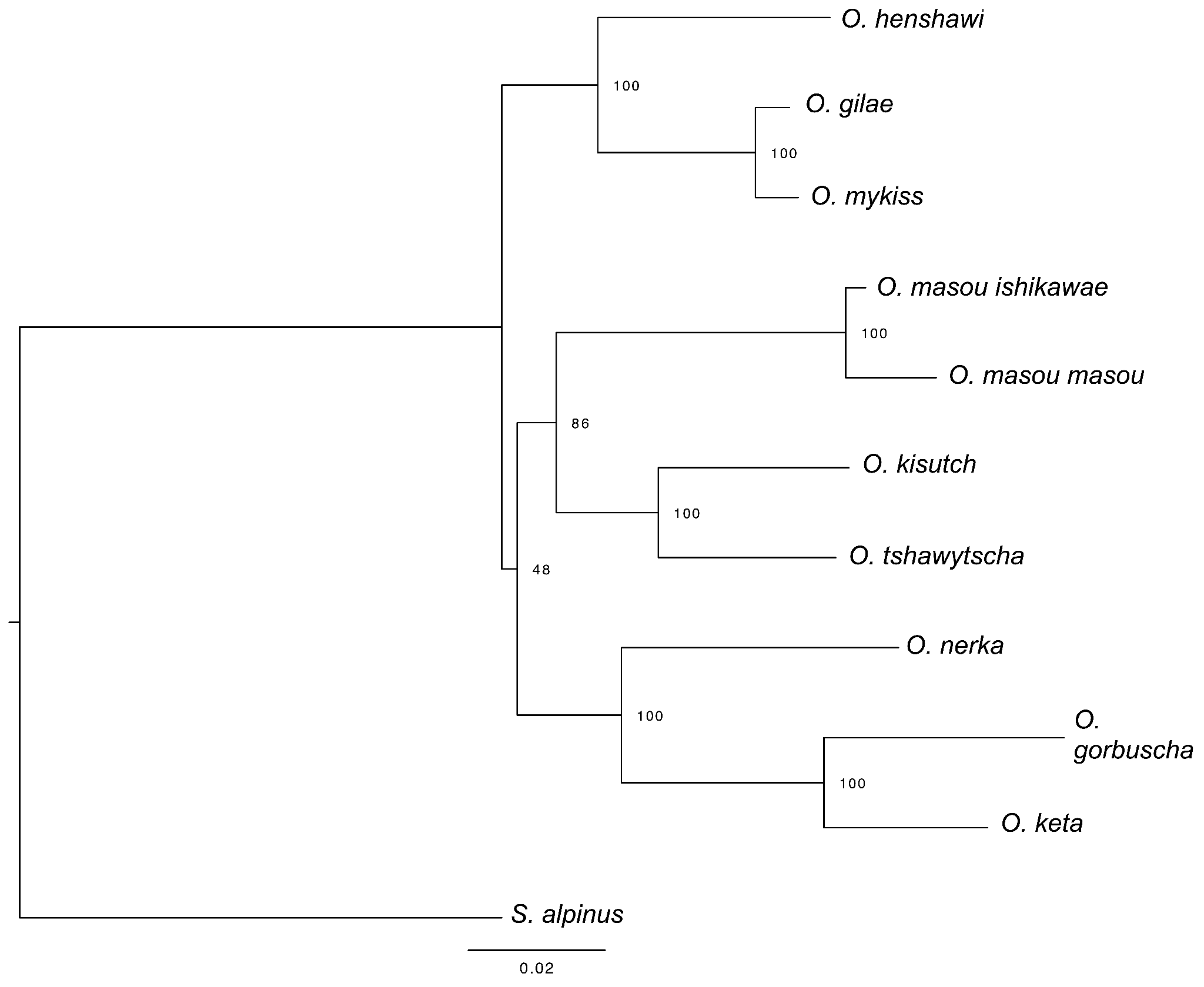

***Supplementary Figure S3. Mitochondrial genome tree of Oncorhynchus.*** *The tree was estimated from the mitochondrial genome alignment in IQ-TREE3. Bootstrap support (1,000 replicates) is shown at each node.*

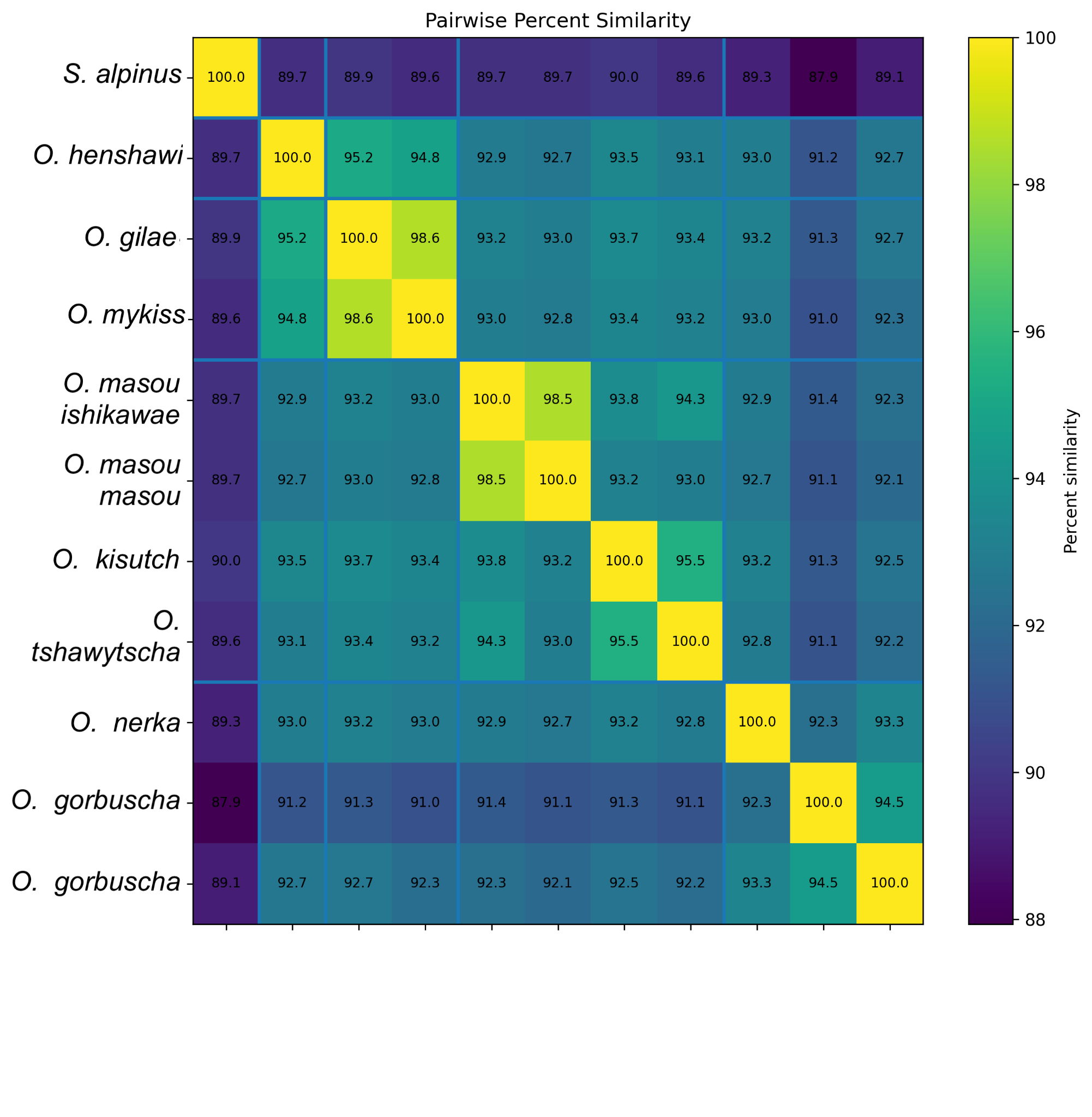

***Supplementary Figure S4.*** *Pairwise mitochondrial genome nucleotide similarity between sampled Oncorhynchus and the outgroup S. alpinus.*

***Supplementary tables***

| ***Alignment ID*** | ***Scientific name*** | ***Common name*** | ***Group in study*** | ***GenBank accession*** | ***Length (bp, RefSeq)*** | ***Un-gapped length in alignment (bp)*** |
| --- | --- | --- | --- | --- | --- | --- |
| *S_alpinus* | *Salvelinus alpinus* | *Arctic char* | *Outgroup* | *NC_000861* | ***~16,655*** | ***16659*** |
| *O_clarkii_henshawi* | *Oncorhynchus henshawi* | *Lahontan cutthroat* | *Trout clade* | *AY886762* | ***~16,675*** | ***16677*** |
| *O_gilae* | *Oncorhynchus gilae* | *Gila trout* | *Trout clade* | *MW300334.1* | ***16,654*** | ***16673*** |
| *O_mykiss_arlee_line* | *Oncorhynchus mykiss* | *Rainbow trout* | *Trout clade* | *L29771 / Arlee assembly GCF_013265735.2* | ***16,660 / assembly-specific*** | ***16642*** |
| *O_masou_ishikawae* | *Oncorhynchus masou ishikawae* | *Amago salmon* | *Not used in group contrast* | *NC_008746* | ***16,658*** | ***16658*** |
| *O_masou_masou* | *Oncorhynchus masou masou* | *Cherry salmon* | *Not used in group contrast* | *NC_008747 / KY320493* | *16,656 / 16,704* | *16652* |
| *O_kisutch* | *Oncorhynchus kisutch* | *Coho salmon* | *Coho / Chinook clade* | *NC_009263* | *~16,659* | *16659* |
| *O_tshawytscha* | *Oncorhynchus tshawytscha* | *Chinook salmon* | *Coho / Chinook clade* | *NC_002980* | *16,644* | *16644* |
| *O_nerka* | *Oncorhynchus nerka* | *Sockeye salmon* | *O. keta + O. gorbuscha + O. nerka clade* | *NC_008615* | *~16,658* | *16658* |
| *O_gorbuscha* | *Oncorhynchus gorbuscha* | *Pink salmon* | *O. keta + O. gorbuscha + O. nerka clade* | *NC_010959 / PP272106* | *~16,679 / ~16,785* | *16785* |
| *O_keta* | *Oncorhynchus keta* | *Chum salmon* | *O. keta + O. gorbuscha + O. nerka clade* | *NC_017838 / KY320492* | *16,652 / 16,656* | *16675* |

***Supplementary Table S1. Mitochondrial genome sequences used in this study.***

| **Donor lineage** | **Recipient lineage** | **Direction** | **NNI** | **Gene trees** | **Sister z** | **Avunc. z** | **djiNNI z** |
| --- | --- | --- | --- | --- | --- | --- | --- |
| O. clarkii lewisi + O. clarkii henshawi + O. mykiss garnerei + O. gilae | *O. kisutch + O. tshawytscha* | unidirectional | 2 | 265 | -8.11 | -5.61 | — |
| *O. tshawytscha* | *O. masou ishikawae + O. masou masou* | bidirectional | 2 | 145 | -3.52 | 15.94 | -2.41 |
| *O. tshawytscha* | *O. masou ishikawae* | bidirectional | 3 | 139 | -11.79 | -10.31 | -7.07 |
| *O. nerka* | *O. kisutch* | bidirectional | 2 | 55 | -3.76 | — | -2.12 |
| *O. masou masou* | *O. nerka + O. keta + O. gorbuscha* | bidirectional | 2 | 26 | -5.1 | — | -3.46 |
| *O. nerka + O. keta + O. gorbuscha* | *O. clarkii henshawi* | unidirectional | 4 | 7 | -2.12 | -1.83 | -1.73 |
| *O. masou ishikawae + O. masou masou* | *O. nerka + O. keta + O. gorbuscha* | — | 1 | 507 | -5.86 | — | — |
| *O. nerka + O. keta + O. gorbuscha* | *O. kisutch* | — | 1 | 148 | -2.77 | — | — |

***Supplementary Table S2.*** *Inferred introgression events (short-branch–corrected run) from the Discordant Attachment Frequency Tool (DAFT). Events are ordered by the number of gene trees showing the discordant attachment. Direction is the inferred direction of gene flow (either unidirectional from the donor to the recipient lineage, or bidirectional between both lineages). The Nearest Neighbor Interchange (NNI) is the number of tree rearrangement operations required for the donor and recipient lineages to be sister in the species tree. Sister z and Avunc. z are the corrected DAFT-test statistics for the pair (values ≤ −1.96 are considered support for introgression). djiNNI z is the bidirectionality-test statistic, reported only where the reverse direction was tested. The final three rows (below the divider) are pairs that are significant but only 1 NNI apart, so djiNNI cannot assign donor versus recipient; their two lineage columns simply list the pair, and their large gene-tree counts partly reflect the close relationship of the lineages.*

*Per-parameter effective sample size (ESS) for each of the four runs, the combined joint ESS across all four runs (chains concatenated), the between-run convergence diagnostic R-hat, and the posterior mean, for four BPP models. R-hat values above 1.1 (poor convergence) are shown in red.*

| Model | Parameter | Post. mean | ESS Run 1 | ESS Run 2 | ESS Run 3 | ESS Run 4 | Joint ESS | R-hat |
| --- | --- | --- | --- | --- | --- | --- | --- | --- |
| masu salmon | theta_9 | 0.0089333 | 34,984 | 16,443 | 53,544 | 38,723 | 143,694 | 1.1324 |
| masu salmon | theta_10 | 0.0062361 | 22,402 | 27,843 | 19,106 | 22,620 | 91,971 | 1.0059 |
| masu salmon | theta_13 | 0.0092326 | 15,016 | 8,232 | 13,142 | 12,798 | 49,187 | 1.0001 |
| masu salmon | theta_14 | 0.0031479 | 30,389 | 22,869 | 28,699 | 30,723 | 112,681 | 1.0035 |
| masu salmon | theta_15 | 0.0034158 | 90,201 | 90,692 | 88,963 | 95,051 | 364,908 | 1.0006 |
| masu salmon | theta_16 | 0.0017742 | 38,898 | 40,689 | 38,210 | 39,096 | 156,894 | 1.0000 |
| masu salmon | theta_17 | 0.0013883 | 128,169 | 126,759 | 132,042 | 128,373 | 515,343 | 1.0000 |
| masu salmon | theta_18 | 0.007031 | 34,108 | 20,498 | 28,632 | 30,349 | 113,587 | 1.0016 |
| masu salmon | theta_19 | 0.0021486 | 9,366 | 6,942 | 8,638 | 8,089 | 33,035 | 1.0645 |
| masu salmon | theta_20 | 0.0010628 | 14,155 | 16,509 | 12,760 | 14,563 | 57,987 | 1.0091 |
| masu salmon | theta_21 | 0.0014037 | 46,213 | 53,717 | 50,353 | 52,671 | 202,954 | 1.0013 |
| masu salmon | theta_22 | 0.013704 | 4,651 | 42,268 | 5,400 | 6,052 | 58,371 | 5.9613 |
| masu salmon | theta_23 | 0.0020208 | 629 | 18,949 | 658 | 703 | 20,939 | 1.7230 |
| masu salmon | tau_13 | 0.011175 | 10,533 | 6,009 | 9,271 | 8,732 | 34,545 | 1.0007 |
| masu salmon | tau_14 | 0.0073734 | 23,479 | 30,670 | 23,535 | 24,903 | 102,587 | 1.0043 |
| masu salmon | tau_15 | 0.0025829 | 47,452 | 44,444 | 46,632 | 48,395 | 186,923 | 1.0000 |
| masu salmon | tau_16 | 0.0017106 | 47,920 | 52,197 | 46,771 | 47,102 | 193,990 | 1.0001 |
| masu salmon | tau_17 | 0.00075702 | 152,246 | 148,388 | 153,357 | 151,702 | 605,694 | 1.0000 |
| masu salmon | tau_18 | 0.0069084 | 8,656 | 5,902 | 7,626 | 7,591 | 29,774 | 1.0346 |
| masu salmon | tau_19 | 0.0064505 | 23,484 | 14,831 | 24,709 | 22,892 | 85,916 | 1.0002 |
| masu salmon | tau_20 | 0.0053398 | 35,787 | 35,426 | 27,598 | 28,737 | 127,548 | 1.0178 |
| masu salmon | tau_21 | 0.0038473 | 66,734 | 75,249 | 66,998 | 71,035 | 280,016 | 1.0004 |
| masu salmon | tau_22 | 0.0017922 | 321 | 58,874 | 329 | 340 | 59,864 | 21.3855 |
| masu salmon | tau_23 | 0.00079943 | 297 | 57,132 | 302 | 309 | 58,040 | 1.9969 |
| masu salmon | M_O_tshawytscha->O_masou_ishikawae | 0.13978 | 24,925 | 26,143 | 25,636 | 27,027 | 103,731 | 1.0013 |
| masu salmon | M_O_masou_ishikawae->O_tshawytscha | 0.011729 | 9,884 | 8,123 | 10,755 | 9,849 | 38,611 | 1.0199 |
| masu salmon | M_tri->masu | 0.036253 | 4,651 | 5,632 | 3,506 | 5,345 | 19,133 | 1.0026 |
| masu salmon | M_masu->tri | 0.0039392 | 2,967 | 3,628 | 2,235 | 2,626 | 11,456 | 1.0175 |
| masu salmon | lnL | -1,245,851.4 | 9,486 | 22,979 | 9,206 | 12,199 | 53,870 | 1.0587 |
| trout salmon introgression | theta_13 | 0.0093214 | 7,932 | 9,175 | 12,353 | 9,828 | 39,288 | 1.0002 |
| trout salmon introgression | theta_14 | 0.0030731 | 11,621 | 14,699 | 18,373 | 14,462 | 59,155 | 1.0002 |
| trout salmon introgression | theta_15 | 0.002714 | 832 | 1,300 | 1,079 | 1,111 | 4,322 | 1.0126 |
| trout salmon introgression | theta_16 | 0.0023867 | 2,528 | 4,278 | 3,504 | 3,920 | 14,230 | 1.0039 |
| trout salmon introgression | theta_17 | 0.0015373 | 5,533 | 9,089 | 7,584 | 7,416 | 29,622 | 1.0018 |
| trout salmon introgression | theta_18 | 0.00765 | 4,996 | 6,618 | 6,163 | 6,647 | 24,423 | 1.0013 |
| trout salmon introgression | theta_19 | 0.00060487 | 589 | 563 | 554 | 556 | 2,262 | 1.0032 |
| trout salmon introgression | theta_20 | 0.0010069 | 5,280 | 6,043 | 6,815 | 6,339 | 24,477 | 1.0011 |
| trout salmon introgression | theta_21 | 0.0014168 | 52,110 | 51,878 | 55,383 | 50,651 | 210,023 | 1.0000 |
| trout salmon introgression | theta_22 | 0.0078851 | 1,276 | 2,120 | 1,708 | 1,954 | 7,059 | 1.0087 |
| trout salmon introgression | theta_23 | 0.0033272 | 144,919 | 124,687 | 154,317 | 146,328 | 570,251 | 1.0001 |
| trout salmon introgression | tau_13 | 0.011147 | 5,235 | 6,519 | 8,148 | 6,897 | 26,799 | 1.0004 |
| trout salmon introgression | tau_14 | 0.0075457 | 5,710 | 6,263 | 7,059 | 6,176 | 25,208 | 1.0004 |
| trout salmon introgression | tau_15 | 0.0029014 | 299 | 344 | 325 | 334 | 1,302 | 1.0472 |
| trout salmon introgression | tau_16 | 0.0016178 | 12,041 | 18,695 | 17,067 | 17,618 | 65,420 | 1.0007 |
| trout salmon introgression | tau_17 | 0.00072763 | 120,800 | 111,228 | 128,875 | 110,068 | 470,970 | 1.0001 |
| trout salmon introgression | tau_18 | 0.0063575 | 1,358 | 2,033 | 1,812 | 2,030 | 7,233 | 1.0069 |
| trout salmon introgression | tau_19 | 0.0062979 | 1,412 | 1,731 | 1,653 | 1,635 | 6,431 | 1.0056 |
| trout salmon introgression | tau_20 | 0.0052982 | 40,166 | 34,402 | 41,665 | 35,830 | 152,063 | 1.0000 |
| trout salmon introgression | tau_21 | 0.003808 | 36,456 | 64,981 | 58,788 | 57,554 | 217,779 | 1.0002 |
| trout salmon introgression | tau_22 | 0.0028826 | 319 | 381 | 353 | 366 | 1,419 | 1.0433 |
| trout salmon introgression | tau_23 | 0.00050295 | 58,476 | 57,627 | 57,912 | 57,954 | 231,969 | 1.0000 |
| trout salmon introgression | M_Trout->Kis_tsh | 0.0070696 | 3,496 | 3,313 | 3,327 | 3,266 | 13,403 | 1.0003 |
| trout salmon introgression | M_Kis_tsh->Trout | 0.013778 | 2,601 | 2,755 | 2,994 | 2,426 | 10,776 | 1.0008 |
| trout salmon introgression | lnL | -1,245,958.9 | 12,002 | 21,097 | 24,398 | 20,887 | 78,385 | 1.0005 |
| within salmon | theta_5 | 0.021524 | 52,299 | 55,769 | 55,920 | 50,387 | 214,375 | 1.0000 |
| within salmon | theta_7 | 0.029854 | 30,317 | 26,721 | 29,933 | 30,863 | 117,835 | 1.0000 |
| within salmon | theta_8 | 0.019627 | 140,718 | 146,477 | 140,013 | 141,871 | 569,078 | 1.0000 |
| within salmon | theta_13 | 0.0092476 | 9,214 | 12,029 | 10,947 | 10,735 | 42,925 | 1.0000 |
| within salmon | theta_14 | 0.0033269 | 51,179 | 67,487 | 72,074 | 60,610 | 251,351 | 1.0000 |
| within salmon | theta_15 | 0.0033928 | 94,549 | 87,104 | 87,182 | 95,152 | 363,986 | 1.0000 |
| within salmon | theta_16 | 0.0017802 | 36,596 | 35,761 | 36,164 | 37,776 | 146,298 | 1.0000 |
| within salmon | theta_17 | 0.0013885 | 125,575 | 122,662 | 123,441 | 125,758 | 497,435 | 1.0000 |
| within salmon | theta_18 | 0.010295 | 9,053 | 14,312 | 10,814 | 12,209 | 46,388 | 1.0001 |
| within salmon | theta_19 | 0.0019599 | 3,016 | 3,712 | 3,257 | 2,904 | 12,889 | 1.0005 |
| within salmon | theta_20 | 0.00056576 | 2,016 | 2,812 | 2,337 | 2,334 | 9,500 | 1.0006 |
| within salmon | theta_21 | 0.0012835 | 7,059 | 5,316 | 6,671 | 5,742 | 24,789 | 1.0003 |
| within salmon | theta_22 | 0.0020621 | 4,789 | 6,236 | 4,884 | 5,045 | 20,954 | 1.0004 |
| within salmon | theta_23 | 0.0032829 | 132,406 | 126,653 | 121,528 | 111,092 | 491,679 | 1.0000 |
| within salmon | tau_13 | 0.011175 | 6,770 | 9,835 | 7,520 | 7,312 | 31,436 | 1.0001 |
| within salmon | tau_14 | 0.0073043 | 66,764 | 74,983 | 83,532 | 81,858 | 307,137 | 1.0000 |
| within salmon | tau_15 | 0.0025829 | 44,447 | 41,232 | 41,024 | 44,645 | 171,349 | 1.0000 |
| within salmon | tau_16 | 0.0017068 | 45,985 | 45,364 | 48,864 | 48,631 | 188,844 | 1.0000 |
| within salmon | tau_17 | 0.00075565 | 153,081 | 154,030 | 150,563 | 153,142 | 610,817 | 1.0000 |
| within salmon | tau_18 | 0.0061339 | 2,999 | 3,981 | 3,194 | 3,769 | 13,943 | 1.0004 |
| within salmon | tau_19 | 0.0060733 | 2,562 | 3,348 | 2,758 | 2,790 | 11,459 | 1.0005 |
| within salmon | tau_20 | 0.0054733 | 3,357 | 5,104 | 4,241 | 4,175 | 16,877 | 1.0003 |
| within salmon | tau_21 | 0.0038227 | 15,031 | 11,724 | 15,833 | 13,655 | 56,243 | 1.0001 |
| within salmon | tau_22 | 0.0047202 | 8,577 | 10,557 | 8,539 | 8,995 | 36,668 | 1.0003 |
| within salmon | tau_23 | 0.00050482 | 52,124 | 51,092 | 49,951 | 51,654 | 204,821 | 1.0000 |
| within salmon | M_O_gorbuscha->O_nerka | 0.053002 | 19,159 | 23,075 | 17,756 | 17,838 | 77,828 | 1.0000 |
| within salmon | M_O_nerka->O_gorbuscha | 0.034885 | 7,292 | 5,830 | 6,567 | 6,176 | 25,864 | 1.0002 |
| within salmon | M_O_nerka->O_kisutch | 0.014822 | 30,583 | 30,827 | 31,981 | 27,604 | 120,995 | 1.0000 |
| within salmon | M_O_kisutch->O_nerka | 0.01607 | 23,725 | 17,446 | 30,764 | 27,968 | 99,904 | 1.0001 |
| within salmon | lnL | -1,246,028.9 | 33,537 | 35,127 | 29,317 | 23,426 | 121,407 | 1.0001 |
| rainbow and cutthroat trout | theta_1 | 0.014466 | 74,976 | 76,855 | 78,369 | 77,144 | 307,344 | 1.0000 |
| rainbow and cutthroat trout | theta_2 | 0.014816 | 95,807 | 99,155 | 99,628 | 100,879 | 395,469 | 1.0000 |
| rainbow and cutthroat trout | theta_4 | 0.012475 | 243,694 | 277,202 | 266,435 | 255,635 | 1,042,966 | 1.0000 |
| rainbow and cutthroat trout | theta_13 | 0.0092438 | 10,035 | 11,638 | 10,697 | 10,030 | 42,399 | 1.0002 |
| rainbow and cutthroat trout | theta_14 | 0.0033243 | 70,962 | 76,567 | 62,123 | 70,413 | 280,065 | 1.0001 |
| rainbow and cutthroat trout | theta_15 | 0.003071 | 83,681 | 83,820 | 82,422 | 88,276 | 338,199 | 1.0000 |
| rainbow and cutthroat trout | theta_16 | 0.0017033 | 45,274 | 46,625 | 45,715 | 44,227 | 181,842 | 1.0000 |
| rainbow and cutthroat trout | theta_17 | 0.0013948 | 78,181 | 83,006 | 78,263 | 81,941 | 321,392 | 1.0000 |
| rainbow and cutthroat trout | theta_18 | 0.010275 | 11,104 | 11,570 | 12,095 | 15,183 | 49,954 | 1.0002 |
| rainbow and cutthroat trout | theta_19 | 0.0013936 | 1,891 | 2,327 | 1,655 | 1,633 | 7,507 | 1.0010 |
| rainbow and cutthroat trout | theta_20 | 0.00070711 | 2,385 | 2,555 | 2,134 | 3,145 | 10,219 | 1.0011 |
| rainbow and cutthroat trout | theta_21 | 0.0014443 | 53,714 | 52,936 | 49,846 | 56,072 | 212,568 | 1.0000 |
| rainbow and cutthroat trout | theta_22 | 0.0022033 | 5,686 | 6,976 | 5,142 | 6,629 | 24,433 | 1.0004 |
| rainbow and cutthroat trout | theta_23 | 0.0032853 | 126,126 | 125,701 | 135,995 | 126,170 | 513,991 | 1.0000 |
| rainbow and cutthroat trout | tau_13 | 0.011175 | 7,881 | 10,341 | 8,087 | 7,744 | 34,053 | 1.0003 |
| rainbow and cutthroat trout | tau_14 | 0.0073064 | 78,990 | 82,022 | 80,498 | 76,982 | 318,492 | 1.0001 |
| rainbow and cutthroat trout | tau_15 | 0.0028605 | 26,396 | 29,101 | 28,933 | 30,008 | 114,438 | 1.0001 |
| rainbow and cutthroat trout | tau_16 | 0.0017444 | 52,366 | 55,223 | 55,029 | 51,787 | 214,404 | 1.0000 |
| rainbow and cutthroat trout | tau_17 | 0.00075658 | 66,669 | 73,543 | 70,248 | 71,834 | 282,294 | 1.0000 |
| rainbow and cutthroat trout | tau_18 | 0.0061498 | 3,782 | 4,241 | 3,651 | 4,592 | 16,265 | 1.0009 |
| rainbow and cutthroat trout | tau_19 | 0.0060863 | 2,723 | 3,353 | 2,505 | 2,915 | 11,496 | 1.0012 |
| rainbow and cutthroat trout | tau_20 | 0.0053755 | 4,767 | 4,756 | 4,177 | 7,776 | 21,476 | 1.0004 |
| rainbow and cutthroat trout | tau_21 | 0.0037832 | 75,473 | 76,341 | 71,402 | 76,287 | 299,504 | 1.0000 |
| rainbow and cutthroat trout | tau_22 | 0.0046917 | 10,351 | 13,884 | 9,281 | 12,340 | 45,856 | 1.0002 |
| rainbow and cutthroat trout | tau_23 | 0.00050474 | 46,648 | 46,026 | 47,698 | 44,943 | 185,315 | 1.0000 |
| rainbow and cutthroat trout | M_O_mykiss_garnerei->O_clarkii_henshawi | 0.047355 | 16,506 | 17,381 | 15,109 | 17,385 | 66,381 | 1.0000 |
| rainbow and cutthroat trout | M_O_mykiss_garnerei->O_clarkii_lewisi | 0.076585 | 10,892 | 12,851 | 12,925 | 12,694 | 49,363 | 1.0001 |
| rainbow and cutthroat trout | M_rainbow_complex->O_clarkii_henshawi | 0.1083 | 21,798 | 22,690 | 22,556 | 23,508 | 90,552 | 1.0001 |
| rainbow and cutthroat trout | M_rainbow_complex->O_clarkii_lewisi | 0.075263 | 10,021 | 10,989 | 11,038 | 11,194 | 43,242 | 1.0002 |
| rainbow and cutthroat trout | lnL | -1,246,025.1 | 15,117 | 36,510 | 50,098 | 11,017 | 112,742 | 1.0006 |

***Supplementary Table S3****. Per-parameter effective sample size (ESS) for each of the four runs, the combined joint ESS across all four runs (chains concatenated), the between-run convergence diagnostic R-hat, and the posterior mean, for four BPP models.*

| **Branch** | **Species tree topology** | | **Mitochondrial genome tree topology** | |
| --- | --- | --- | --- | --- |
|  | **Mean** | **95% HPD** | **Mean** | **95% HPD** |
| *Oncorhynchus* stem | 4.68 | 1.62–8.60 | 4.61 | 1.39–8.75 |
| Pacific trout stem | 2.78 | 1.53–4.26 | 2.55 | 1.31–4.04 |
| *O. henshawi* | 4.17 | 3.20–5.48 | 4.12 | 3.17–5.48 |
| *O. gilae* + *O. mykiss* stem | 3.58 | 2.49–5.04 | 3.61 | 2.49–5.18 |
| *O. gilae* | 3.19 | 1.80–4.60 | 3.10 | 1.68–4.52 |
| *O. mykiss* | 3.83 | 2.17–5.48 | 3.74 | 2.10–5.47 |
| Pacific salmon stem | 4.01 | 1.44–7.09 | 3.83 | 1.22–7.04 |
| *O. kisutch* + *O. tshawytscha* + *O. keta* + *O. gorbuscha* + *O. nerka* stem | 3.81 | 1.22–6.77 | – | – |
| *O. kisutch* + *O. tshawytscha* + *O. masou* stem | – | – | 3.74 | 1.27–6.63 |
| *O. keta* + *O. gorbuscha* + *O. nerka* stem | 5.46 | 2.88–8.66 | 4.72 | 2.01–7.79 |
| *O. gorbuscha* + *O. keta* stem | 6.91 | 4.16–10.32 | 6.69 | 3.97–10.12 |
| *O. gorbuscha* | 6.35 | 4.51–8.33 | 6.19 | 4.30–8.22 |
| *O. keta* | 4.46 | 3.12–5.88 | 4.32 | 2.99–5.78 |
| *O. nerka* | 4.10 | 3.30–5.02 | 3.96 | 3.19–4.84 |
| *O. kisutch* + *O. tshawytscha* stem | 3.67 | 2.07–5.76 | 3.80 | 1.79–6.30 |
| *O. kisutch* | 3.62 | 2.56–4.77 | 3.71 | 2.60–4.87 |
| *O. tshawytscha* | 3.45 | 2.40–4.53 | 3.41 | 2.39–4.50 |
| *O. masou* stem | 4.11 | 3.32–5.03 | 4.47 | 3.21–5.84 |
| *O. masou ishikawae* | 2.06 | 1.17–2.92 | 1.65 | 0.88–2.43 |
| *O. masou masou* | 4.75 | 2.98–6.33 | 5.16 | 3.20–6.85 |
| *S. alpinus* | 6.21 | 3.65–8.84 | 6.27 | 3.64–8.82 |

***Supplementary Table S4****. Branch-specific substitution rates estimated in MCMCtree on the species tree and mitochondrial genome tree topologies. Values are posterior means and 95% highest posterior density (HPD) intervals in units of 10^−3^ substitutions/site/Myr, from two independent chains merged after burn-in. Stem branches are named by the clade they subtend; dashes indicate branches present in only one topology. Because the trees were rooted with* S. alpinus*, the outgroup branch and the* Oncorhynchus *stem are estimated separately.*

| **Branch / statistic** | **Concat.** | **ND1** | **ND2** | **COI** | **COII** | **ATP8** | **ATP6** | **COIII** | **ND3** | **ND4L** | **ND4** | **ND5** | **ND6** | **CYTB** |
| --- | --- | --- | --- | --- | --- | --- | --- | --- | --- | --- | --- | --- | --- | --- |
| Codons | 3801 | 324 | 349 | 516 | 230 | 55 | 227 | 261 | 116 | 98 | 460 | 612 | 173 | 380 |
| **Species tree topology** | | | | | | | | | | | | | | |
| *S. alpinus* + *Oncorhynchus* stem | 0.026 | 0.009 | 0.055 | 0.007 | 0.017 | 0.182 | 0.008 | 0.039 | 0.061 | 0.000 | 0.033 | 0.038 | 0.017 | 0.007 |
| Pacific trout stem | 0.021 | 0.000 | 0.093 | 0.000 | 0.000 | 0.000 | 0.000 | 0.000 | 0.000 | 0.000 | 0.000 | 0.051 | 0.000 | 0.000 |
| *O. henshawi* | 0.028 | 0.034 | 0.049 | 0.000 | 0.020 | 0.000 | 0.000 | 0.000 | 0.000 | 0.000 | 0.038 | 0.053 | 0.015 | 0.015 |
| *O. gilae* + *O. mykiss* stem | 0.019 | 0.000 | 0.049 | 0.000 | 0.000 | 0.000 | 0.000 | 0.026 | 0.028 | 0.000 | 0.032 | 0.043 | 0.000 | 0.000 |
| *O. gilae* | 0.056 | 0.000 | 0.152 | 0.000 | – | 0.000 | 0.000 | 0.000 | 0.000 | – | 0.186 | 0.000 | – | 0.000 |
| *O. mykiss* | 0.125 | 0.148 | 0.235 | 0.128 | 0.137 | 0.000 | 0.000 | 0.000 | 0.000 | – | 0.037 | 0.439 | 0.000 | 0.094 |
| Pacific salmon stem | 0.028 | 0.000 | 0.000 | 0.000 | 0.258 | – | – | 0.135 | 0.000 | 0.000 | 0.000 | 0.064 | 0.000 | 0.000 |
| *O. kisutch* + *O. tshawytscha* + *O. keta* + *O. gorbuscha* + *O. nerka* stem | – | 0.000 | – | – | – | – | – | – | – | – | 0.092 | – | – | – |
| *O. keta* + *O. gorbuscha* + *O. nerka* stem | 0.017 | 0.000 | 0.018 | 0.056 | 0.083 | – | 0.000 | 0.026 | 0.000 | 0.000 | 0.000 | 0.027 | 0.000 | 0.000 |
| *O. gorbuscha* + *O. keta* stem | 0.014 | 0.015 | 0.025 | 0.015 | 0.000 | – | 0.015 | 0.000 | 0.000 | 0.000 | 0.009 | 0.010 | 0.041 | 0.000 |
| *O. gorbuscha* | 0.017 | 0.000 | 0.009 | 0.038 | 0.000 | 0.000 | 0.000 | 0.000 | 0.000 | 0.000 | 0.016 | 0.021 | 0.076 | 0.024 |
| *O. keta* | 0.018 | 0.012 | 0.039 | 0.000 | 0.000 | 0.000 | 0.000 | 0.024 | 0.016 | 0.000 | 0.010 | 0.033 | 0.159 | 0.000 |
| *O. nerka* | 0.025 | 0.018 | 0.026 | 0.000 | 0.048 | 0.000 | 0.020 | 0.000 | 0.076 | 0.000 | 0.023 | 0.047 | 0.018 | 0.000 |
| *O. kisutch* + *O. tshawytscha* stem | 0.008 | 0.000 | 0.013 | 0.000 | 0.000 | 0.000 | 0.000 | 0.000 | – | 0.000 | 0.022 | 0.026 | 0.000 | 0.000 |
| *O. kisutch* | 0.029 | 0.087 | 0.000 | 0.000 | 0.045 | 0.000 | 0.000 | 0.000 | 0.048 | 0.000 | 0.072 | 0.048 | 0.000 | 0.010 |
| *O. tshawytscha* | 0.034 | 0.013 | 0.093 | 0.000 | 0.031 | – | 0.020 | 0.000 | 0.191 | 0.000 | 0.040 | 0.018 | 0.077 | 0.030 |
| *O. masou* stem | 0.028 | 0.008 | 0.089 | 0.000 | 0.000 | 0.000 | 0.015 | 0.000 | 0.000 | 0.028 | 0.035 | 0.037 | 0.053 | 0.029 |
| *O. masou ishikawae* | 0.055 | 0.080 | 0.411 | 0.000 | 0.000 | – | 0.000 | 0.000 | – | – | 0.000 | 0.034 | 1.312 | 0.000 |
| *O. masou masou* | 0.056 | 0.018 | – | 0.000 | – | – | 0.000 | 0.000 | 0.027 | – | 0.025 | 0.106 | 0.090 | 0.059 |
| One-ratio (M0) ω | 0.026 | 0.016 | 0.050 | 0.007 | 0.025 | 0.027 | 0.007 | 0.014 | 0.038 | 0.003 | 0.028 | 0.040 | 0.036 | 0.011 |
| lnL, M0 | -36124.10 | -3260.27 | -3625.62 | -4406.71 | -1725.35 | -351.33 | -2128.84 | -2169.83 | -1218.22 | -769.17 | -4675.48 | -5965.99 | -1751.72 | -3603.62 |
| lnL, free-ratio | -36093.29 | -3246.31 | -3606.42 | -4392.82 | -1714.85 | -347.94 | -2125.19 | -2159.45 | -1208.01 | -767.14 | -4664.30 | -5954.71 | -1736.85 | -3588.50 |
| 2ΔlnL (df = 18) | 61.61 | 27.93 | 38.41 | 27.79 | 20.99 | 6.77 | 7.30 | 20.76 | 20.43 | 4.06 | 22.38 | 22.55 | 29.73 | 30.24 |
| ΔAIC (free-ratio − M0) | -25.6 | 8.1 | -2.4 | 8.2 | 15.0 | 29.2 | 28.7 | 15.2 | 15.6 | 31.9 | 13.6 | 13.4 | 6.3 | 5.8 |
| *P* | <0.001 | 0.063 | 0.003 | 0.065 | 0.280 | 0.992 | 0.987 | 0.292 | 0.309 | 1.000 | 0.216 | 0.208 | 0.040 | 0.035 |
| *P* (Holm) | – | 0.631 | **0.044** | 0.631 | 1.000 | 1.000 | 1.000 | 1.000 | 1.000 | 1.000 | 1.000 | 1.000 | 0.441 | 0.422 |
| **Mitochondrial genome tree topology** | | | | | | | | | | | | | | |
| *S. alpinus* + *Oncorhynchus* stem | 0.026 | 0.009 | 0.058 | 0.007 | 0.017 | 0.182 | 0.008 | 0.037 | 0.065 | 0.000 | 0.028 | 0.038 | 0.017 | 0.007 |
| Pacific trout stem | 0.022 | 0.000 | 0.076 | 0.000 | 0.000 | 0.000 | 0.000 | 0.000 | 0.000 | 0.000 | 0.024 | 0.057 | 0.000 | 0.000 |
| *O. henshawi* | 0.028 | 0.034 | 0.043 | 0.000 | 0.020 | 0.000 | 0.000 | 0.000 | 0.000 | 0.000 | 0.044 | 0.053 | 0.015 | 0.016 |
| *O. gilae* + *O. mykiss* stem | 0.018 | 0.000 | 0.049 | 0.000 | 0.000 | 0.000 | 0.000 | 0.025 | 0.029 | 0.000 | 0.022 | 0.042 | 0.000 | 0.000 |
| *O. gilae* | 0.058 | 0.000 | 0.168 | 0.000 | – | 0.000 | 0.000 | 0.000 | 0.000 | – | 0.187 | 0.000 | – | 0.000 |
| *O. mykiss* | 0.122 | 0.147 | 0.210 | 0.127 | 0.138 | 0.000 | 0.000 | 0.000 | 0.000 | – | 0.037 | 0.464 | 0.000 | 0.091 |
| Pacific salmon stem | 0.044 | 0.000 | 0.000 | 0.000 | 0.260 | – | – | 0.152 | – | 0.000 | 0.000 | 0.102 | 0.000 | – |
| *O. kisutch* + *O. tshawytscha* + *O. masou* stem | 0.002 | 0.000 | – | 0.000 | – | – | 0.000 | 0.000 | 0.000 | – | 0.026 | 0.000 | 0.000 | 0.000 |
| *O. keta* + *O. gorbuscha* + *O. nerka* stem | 0.018 | 0.000 | 0.000 | 0.056 | 0.167 | – | 0.000 | 0.029 | 0.000 | 0.000 | 0.000 | 0.032 | 0.000 | 0.000 |
| *O. gorbuscha* + *O. keta* stem | 0.014 | 0.015 | 0.024 | 0.015 | 0.000 | – | 0.015 | 0.000 | 0.000 | 0.000 | 0.009 | 0.010 | 0.041 | 0.000 |
| *O. gorbuscha* | 0.017 | 0.000 | 0.009 | 0.038 | 0.000 | 0.000 | 0.000 | 0.000 | 0.000 | 0.000 | 0.016 | 0.021 | 0.077 | 0.025 |
| *O. keta* | 0.018 | 0.012 | 0.039 | 0.000 | 0.000 | 0.000 | 0.000 | 0.024 | 0.017 | 0.000 | 0.010 | 0.034 | 0.153 | 0.000 |
| *O. nerka* | 0.025 | 0.018 | 0.026 | 0.000 | 0.048 | 0.000 | 0.020 | 0.000 | 0.084 | 0.000 | 0.018 | 0.048 | 0.018 | 0.000 |
| *O. kisutch* + *O. tshawytscha* stem | 0.010 | 0.000 | 0.000 | 0.000 | 0.031 | 0.000 | 0.000 | 0.000 | – | 0.000 | 0.032 | 0.035 | 0.000 | 0.000 |
| *O. kisutch* | 0.029 | 0.082 | 0.000 | 0.000 | 0.045 | 0.000 | 0.000 | 0.000 | 0.046 | 0.000 | 0.069 | 0.047 | 0.000 | 0.009 |
| *O. tshawytscha* | 0.034 | 0.014 | 0.093 | 0.000 | 0.031 | – | 0.020 | 0.000 | 0.240 | 0.000 | 0.041 | 0.019 | 0.080 | 0.036 |
| *O. masou* stem | 0.031 | 0.008 | 0.089 | 0.000 | 0.000 | 0.000 | 0.016 | 0.000 | 0.000 | 0.028 | 0.042 | 0.038 | 0.057 | 0.032 |
| *O. masou ishikawae* | 0.076 | 0.113 | 0.411 | 0.000 | 0.000 | – | 0.000 | – | – | – | 0.000 | 0.066 | – | 0.000 |
| *O. masou masou* | 0.049 | 0.016 | – | 0.000 | – | – | 0.000 | 0.000 | 0.027 | – | 0.022 | 0.085 | 0.078 | 0.047 |
| One-ratio (M0) ω | 0.026 | 0.016 | 0.050 | 0.007 | 0.027 | 0.027 | 0.007 | 0.014 | 0.038 | 0.003 | 0.028 | 0.041 | 0.036 | 0.011 |
| lnL, M0 | -36106.83 | -3257.32 | -3625.37 | -4406.65 | -1729.77 | -351.33 | -2127.64 | -2166.11 | -1209.57 | -769.17 | -4670.00 | -5962.39 | -1751.29 | -3595.09 |
| lnL, free-ratio | -36077.34 | -3243.22 | -3603.30 | -4392.77 | -1720.54 | -347.94 | -2123.95 | -2156.07 | -1197.98 | -767.14 | -4659.65 | -5950.04 | -1735.33 | -3581.20 |
| 2ΔlnL (df = 18) | 58.98 | 28.20 | 44.14 | 27.77 | 18.48 | 6.77 | 7.38 | 20.10 | 23.18 | 4.06 | 20.72 | 24.71 | 31.92 | 27.79 |
| ΔAIC (free-ratio − M0) | -23.0 | 7.8 | -8.1 | 8.2 | 17.5 | 29.2 | 28.6 | 15.9 | 12.8 | 31.9 | 15.3 | 11.3 | 4.1 | 8.2 |
| *P* | <0.001 | 0.059 | <0.001 | 0.066 | 0.425 | 0.992 | 0.987 | 0.327 | 0.184 | 1.000 | 0.294 | 0.133 | 0.022 | 0.065 |
| *P* (Holm) | – | 0.651 | **0.007** | 0.653 | 1.000 | 1.000 | 1.000 | 1.000 | 1.000 | 1.000 | 1.000 | 1.000 | 0.270 | 0.653 |

***Supplementary Table S5****. Gene-wise ratios of nonsynonymous to synonymous substitution rates (dN/dS, ω) estimated in codeml under the free-ratio (one ω per branch) and one-ratio (M0; one ω across the tree) models for the concatenated mitochondrial protein-coding alignment (Concat.) and each of the 13 genes, on the species tree and mitochondrial genome tree topologies. Branch values are free-ratio maximum-likelihood estimates; stem branches are named by the clade they subtend. Because codeml models are unrooted, ω for the* S. alpinus *branch and the* Oncorhynchus *stem is estimated jointly. Dashes indicate branches on which no synonymous change was inferred (dS = 0), for which ω is undefined. Models were compared with likelihood-ratio tests (2ΔlnL, df = 18) and the Akaike Information Criterion (ΔAIC < 0 favors the free-ratio model); gene-wise* P*-values were Holm-corrected across the 13 genes within each topology, and Holm-adjusted values in bold are significant (*P *< 0.05). Each model was run from at least two starting values that converged on the same maximum log-likelihood (Supplementary Note S6).*

**Supplementary Note S1: divergence time estimation in BEAST2**

As inter-lineage nucleotide substitution rate variation can cause the false-positive signal for introgression (Frankel and Ané 2023), we used BEAST v2.7.7 (Bouckaert et al. 2014, 2019) to estimate both divergence times and lineage-specific substitution rates across our 12 taxa, allowing us to assess whether rate heterogeneity could account for any introgression we recovered. Divergence times are also necessary for evaluating whether hybridization between lineages was temporally feasible. We filtered for loci concordant with the species tree topology, selected 100 loci with the highest average bootstrap support (>68.5% average ultra-fast bootstrap support) and then concatenated the alignments into a single concatenated supermatrix with FASconCAT v1.11 (Kück and Meusemann 2010). We utilized the Generalized Time Reversible nucleotide substitution model (Tavaré 1986) with four gamma rate categories (Yang 1994), estimating the mean substitution rate. We employed an optimized relaxed clock (Drummond et al. 2006; Zhang and Drummond 2020), allowing rates to vary across the tree and between loci. Tree shape and divergence times were modeled under the birth–death prior (Gernhard 2008; Douglas et al. 2021).

We used uniform distributions for all time constraints, with the same fossil calibrations as Lecaudey et al. (2018), who generated a time-tree of the broader Salmoninae with RADseq and included eight of the *Oncorhynchus* species considered here. We set the maximum root age of *Oncorhynchus* and *Salvelinus* to 25 Ma, based upon the upper estimates recovered by Lecaudey et al. (2018). *Oncorhynchus rastrosus* is the oldest known representative of *Oncorhynchus*, at 11.5 Ma (±0.5 Ma) (Eiting and Smith 2007) and a member of the Pacific salmon clade, thus we set 11 Ma as the minimum age constraint for the crown node of this clade. The species *Oncorhynchus ketopsis*, dated to between 6 and 8 Ma in the late Miocene, is considered to belong to the same clade as *O. gorbuscha*, *O. keta*, and *O. nerka* (Eiting and Smith 2007), and thus was used to set the minimum age of these three taxa to 6 Ma. As the oldest Pacific trout, we used *O. lacustris* as justification for setting the minimum age of the Pacific trout to 3.2 Ma (Crête-Lafrenière et al. 2012). All other priors were implemented with the default values assigned by BEAUti v2.7.7 (Bouckaert et al. 2014, 2019). We conducted four independent Markov Chain Monte Carlo (MCMC) runs for 100 million generations, sampling trees and logging parameters every 1000th generation. We dropped the first 25% of samples as burn-in and assessed convergence of the four independent runs using TRACER (Rambaut et al. 2018), with a joint ESS > 200 across model parameters considered as the minimum cutoff for convergence. We then combined the post-burn-in trees from each run and estimated the maximum clade credibility tree using TreeAnnotator v2.7.7 (Bouckaert et al. 2014).

**Supplementary Note S2: applying the PSMC for comparison with BEAST divergence time estimations**

As hatchery individuals are bred to have high homozygosity, we only used the genome assemblies from wild individuals, thus *O. mykiss*, *O. kisutch*, and *O. tshawytscha* were excluded from this analysis, as was *O. gilae*, for which sequencing reads were not available at the time of analysis. We mapped the reads from the Pacific salmon to the reference assembly of *O. gorbuscha*, and from the Pacific trout to *O. henshawi*. For the assemblies of *O. gorbuscha* and *O. henshawi*, we mapped ONT reads to their respective genome assemblies with minimap2 v2.1 (Li 2018) using the flag map-ont. We also used minimap2 v2.1 (Li 2018) to map the PacBio HiFi reads from *O. nerka* to the *O. gorbuscha* reference assembly, with the flag asm20. As long reads were only used for scaffolding the genome assemblies of *O. masou masou*, *O. masou ishikawae*, and *O. lewisi*, we mapped the Illumina reads for each species to the reference assemblies with BWA v0.7.17-r1188 (Li and Durbin 2009). We compressed each of the resulting SAM files into sorted and indexed BAM files with samtools v1.16.1 (Danecek et al. 2021), and then called bases from sites with a minimum coverage depth of 15, and a maximum depth of 125 (these parameters were shifted to 15 and 135 for *O. henshawi*, and 15 and 200 for *O. gorbuscha* to account for the higher average read coverage of these assemblies) with samtools v1.16.1 and bcftools v1.6 (Danecek et al. 2021), employing default filtering settings. We used psmc v0.6.5-r67 (Li 2023) to convert the base-calling output to psmcfa format and ran each PSMC analysis with 100 bootstrap replicates. To avoid false positive peaks in effective population size (N_e_) we split the first time window in the model into two windows (Hilgers et al. 2025). We visualized the resulting demographic models with gnuplot v5.2 (Phillips 2012). For each species, we assumed a genome-wide mutation rate of 3.0e-9, as has previously been used in salmonids (Subramanian and Kumar 2023). We used a generation time of four years for both masu salmon (Ueda 2018), four and a half years for *O. nerka* (National Marine Fisheries Service 2022), two years for *O. gorbuscha* (Christensen et al. 2021), and five years for both *O. lewisi* (Muhlfeld et al. 2012) and *O. henshawi* (Montana Natural Heritage Program n.d.).

**Supplementary Note S3: introgression mapping with GhostParser and species network inference with PhyloNet**

We used GhostParser (Tolman and Suvorov 2025) and the Discordant Attachment Frequency Tool (DAFT; Mishra et al. 2026) to map introgression events onto our estimated species tree. We first used GhostParser v1.0 to test for evidence of both sampled and ghost (involving no sampled lineages) introgression in our tree. GhostParser first employs the Discordant Counts Test (DCT; Suvorov et al. 2022a,b), which tests for elevated counts of one discordant (not matching the species tree) topology relative to the other across triplets. For instances where a signal of introgression is found, GhostParser compares the heights of the concordant and most abundant discordant topologies to infer the recipient of introgression and whether the donor is a sampled or ghost lineage. We used the coalescent species tree as the guide tree, the bootstrapped gene trees as input, and S. alpinus as the outgroup. We used a chi-square test (P = 0.01, to retain only highly supported putative introgression events) to implement the DCT, and a KS test (P = 0.05) to compare the median heights of gene trees, calculated as the average of all possible pathways.

We inferred the species network using the pseudo-likelihood framework from InferNetwork_MPL (MPL; Yu and Nakhleh 2015; Cao and Nakhleh 2019) as implemented in PhyloNet v0.3.0 (Than et al. 2008), with all of our estimated gene trees as input. We tested maximum numbers of allowed reticulations from 0 to 4, and replicated each run four times, for a total of 16 independent runs. To select the best network we pooled the top five networks from each independent run and, for each, calculated the Akaike Information Criterion (AIC; Akaike 1974) and Bayesian Information Criterion (BIC; Schwarz 1978) with a custom Python script (Python v3.10.12), as AIC = 2K − 2lnL and BIC = K·ln(n) − 2lnL, where lnL is the (pseudo-)log-likelihood reported by PhyloNet, n is the number of gene trees analyzed (2,486), and K is the number of free parameters of the network (its estimated internal branch lengths plus inheritance probabilities, K = t + 4r − 2 for a network with t taxa and r reticulations) (Yu et al. 2014). We ranked networks by AIC, with the lowest value indicating the preferred model, as implemented for phylogenetic-network model selection in PhyloNet (Wen et al. 2018).

**Supplementary Note S4: multispecies-coalescent-with-migration models in BPP**

To further test the support for the putative introgression events with GhostParser, DAFT, and MPL (PhyloNet), we fit four Multispecies-Coalescent-with-Migration (MSC-M) models in Bayesian Phylogenetics and Phylogeography (BPP) v4.7.0 (Flouri et al. 2020, 2023) to our locus alignments. The MSC-M model assumes that gene flow is continuous over extended periods of time (Hey et al. 2018; Flouri et al. 2023) and is highly sensitive to low levels of introgression (Pang and Zhang 2024; Ji et al. 2025). Models 1–3 tested candidate events from GhostParser, DAFT and MPL, while model 4 tested gene flow between rainbow and cutthroat trout lineages. The first model tested the introgression identified between *O. gorbuscha* and *O. nerka*, and between *O. nerka* and *O. kisutch*, both supported by DAFT. The second included migration between *O. masou ishikawae* and *O. tshawytscha* (identified by GhostParser and DAFT), and between the MRCA of masu salmon, and the ancestor of *O. gorbuscha*, *O. keta*, and *O. nerka* (supported by DAFT). The third model tested migration between the ancestor of Pacific trout and the ancestor of *O. kisutch and O. tshawytscha*, which would result in introgression supported by all three of the DAFT, GhostParser, and MPL techniques. Although none of the preceding analyses detected gene flow between rainbow and cutthroat trout lineages, we fit a fourth model to test for it explicitly, given its conservation importance. This model contained only migration between the ancestor of O. gilae + O. mykiss and each of O. lewisi and O. henshawi, and between O. mykiss and each of O. lewisi and O. henshawi. We specified all migration events as bidirectional. We implemented BPP with the same model parameters as Ji et al. (2025). This included assigning the migration rate to the gamma prior *M ~* G(2, 10), with mean 0.2. We specified a burn-in of 500,000 iterations and then took one million samples, sampling every 2 iterations. We conducted each run twice, and checked for convergence between model parameters at ESS > 200, visual inspection of trace plots, and the Gelman-Rubin statistic (runs were considered converged at R̂ < 1.1). To determine whether the model supported the retention of each individual migration parameter, we calculated Bayes Factors (*B10*) for each migration parameter via the Savage-Dickey Ratio (Ji et al. 2023, 2025), where M is the population migration rate (expected number of migrants per generation) and the null hypothesis of no gene flow is approximated as M < ε, with ε a rate treated as effectively zero. Because B₁₀ is sensitive to this choice, we used both ε = 0.001 and ε = 0.0001. We retained gene flow events with *B10* > 100 in the final models, which is considered decisive (Kass and Raftery 1995).

**Supplementary Note S5: applying the PSMC to detect introgression from a hybrid genome**

To use the hybrid *O. mykiss* × *O. lewisi* assembly to test for deep historical introgression between *O. lewisi* and *O. mykiss*, we used the PSMC as a means of identifying coalescence between the two parental lineages. In typical diploid individuals that are not F1 hybrids, the PSMC is used to infer historical changes in effective population size (N_e_) over time. It does so by modeling the distribution of coalescent events between two alleles at loci across the genome, estimating how frequently alleles coalesce within specific time intervals in the past (Mather et al. 2020). The PSMC employs a hidden Markov model, in which the time to the most recent common ancestor (MRCA) of alleles is treated as a hidden state underlying the observed sequence data (Mather et al. 2020). Because N_e_ is inversely correlated with the rate of coalescence, short intervals between coalescent events imply a small N_e_, while longer intervals suggest a larger N_e_. Under coalescent theory, the inverse of the coalescence rate serves as a proxy for N_e_, assuming the absence of confounding factors such as natural selection or nonrandom mating (Mazet et al. 2016; Mather et al. 2020). In an F1 hybrid whose parental lineages were completely isolated between speciation and hybridization, all alleles are expected to coalesce after the speciation event, when tracing the two lineages backwards in time (Douglas et al. 2014; Cahill et al. 2016; Beichman et al. 2018). Therefore, any coalescence more recent than the divergence of the parental lineages, which cannot arise from incomplete lineage sorting, would provide evidence of post-speciation gene flow between them, provided the underlying assumptions of the PSMC (e.g., no selection on alleles, no rate variation between alleles; Mather et al. 2020) are met. An absence of coalescence prior to speciation indicates that no hybridization between the lineages has occurred, or that all introgressed loci have been purged from the genomes of *both* parental lineages (Joly et al. 2009; Veller et al. 2023).

**Supplementary Note S6: mitochondrial divergence times, substitution rates, and dN/dS**

To estimate lineage-specific substitution rates in the mitochondrial genome, we dated the mitochondrial genome alignment (16,858 columns) on two fixed topologies rooted with *Salvelinus alpinus*. The first was the nuclear species tree inferred in BEAST v2.7.7 (Bouckaert et al. 2019), pruned to the taxa in the mitochondrial dataset. The second was the maximum-likelihood mitochondrial genome tree inferred in IQ-TREE v3.1.1 (Wong et al. 2025). Divergence times and branch-specific substitution rates were estimated in MCMCtree in PAML v4.10.10 (Yang 2007) using the approximate likelihood method (dos Reis and Yang 2011), with the gradient and Hessian of branch lengths calculated in baseml under GTR+Γ₅ (Tavaré 1986; Yang 1994). Rates were modelled with an independent log-normal relaxed clock (Rannala and Yang 2007), using gamma priors on the mean rate (α = 2, β = 4; mean 0.5 substitutions/site/100 Myr) and on the rate variance (α = 1, β = 10), and a birth–death tree prior (λ = μ = 1, ρ = 0.1; multiplicative construction). Node ages were constrained with soft-bounded uniform calibrations (2.5% tail probabilities; Yang and Rannala 2006) matching the 95% highest posterior density (HPD) intervals of node heights in the BEAST species tree. All ten internal nodes were calibrated on the species topology. On the mitochondrial topology, only nodes shared with the species tree were calibrated, leaving the most recent common ancestor of Pacific salmon and the node uniting *O. kisutch* + *O. tshawytscha* with masu salmon unconstrained. For each topology, we ran two independent chains, each with a burn-in of 2 × 10⁴ iterations followed by 2 × 10⁴ samples drawn every 100 iterations. Post-burn-in samples from both chains were combined, and convergence was assessed with the effective sample size (ESS), summed across chains, and R̂.

To evaluate selective constraint, we extracted the 13 mitochondrial protein-coding genes using the gene coordinates of the annotated *O. gilae* reference mitogenome. Each gene was aligned in a codon-aware manner in Biopython v1.88 (Cock et al. 2009). Sequences were translated under the vertebrate mitochondrial genetic code and aligned at the amino acid level to the longest sequence using the BLOSUM62 matrix (Henikoff and Henikoff 1992). The original nucleotide sequences were then threaded onto the amino acid alignment. Terminal stop codons, including incomplete stop codons completed by polyadenylation, were removed, ND6 was reverse-complemented, and codons between compensating frameshifts in ND5 of *O. mykiss* and *O. tshawytscha* were coded as missing data, yielding a concatenated alignment of 3,801 codons. Branch-specific ratios of nonsynonymous to synonymous substitution rates (dN/dS, ω) were estimated with codeml in PAML v4.10.10 under the codon substitution model of Goldman and Yang (1994) with F3x4 codon frequencies. We fitted the free-ratio model (Yang 1998) on both topologies; because codeml models are unrooted, ω on the outgroup branch and the *Oncorhynchus* stem was estimated jointly. To test whether ω varied among branches, the free-ratio model was compared with the one-ratio model (M0), in which a single ω is shared by all branches, for the concatenated alignment and for each of the 13 genes separately, on both topologies. Twice the log-likelihood difference (2ΔlnL) was compared with a χ² distribution with 18 degrees of freedom (19 branch-specific ω parameters versus one), and gene-wise *P*-values were corrected for multiple testing within each topology with the Holm (1979) method. Each model was fitted twice, from initial values of ω = 0.05 and κ = 2, and ω = 0.8 and κ = 8. All M0 fits and 22 of the 28 free-ratio fits reached the same log-likelihood (within 0.01 units) from both starts. For the remaining six free-ratio fits, additional runs were started from ω = 2 and κ = 1; ω = 0.2 and κ = 4; and ω = 0.01 and κ = 15 (and, for COII on the mitochondrial topology, from four further starting values), until at least two runs reached the same highest log-likelihood; estimates from the highest-likelihood run are reported (Table S5). Control files, tree files, and output for all codeml and MCMCtree analyses are provided as Supplementary Data.
